# Structural basis of the protein kinase PKN1 HR1 domain oligomerization and differential regulation by RhoA and Rac1

**DOI:** 10.64898/2026.01.30.702881

**Authors:** Georgios Sophocleous, Darerca Owen, Helen R. Mott

## Abstract

The protein kinase C-related kinase (PKN) family of serine/threonine kinases consists of PKN1, PKN2 and PKN3, all of which are Rho family GTPase effectors. PKNs have three N-terminal Homology Region 1 (HR1) domains (HR1a, HR1b and HR1c), which form antiparallel coiled coils, which in two cases interact with Rho family GTPases, activating the kinase. The PKNs are implicated in several important cellular processes, including cytoskeletal regulation, cell adhesion, gene expression and cell cycle progression, and are also implicated in cancer. Here we have investigated the roles of the HR1 domains in PKN oligomerisation. We show that PKN1 HR1a is a dimer and that the HR1c domain drives further oligomerization. We have mapped the interactions between the HR1 domains and used an integrative approach to model HR1-containing PKN1 dimers. Biophysical analysis shows that RhoA forms a 1:2 complex with HR1a, resulting in a rearrangement of the HR1a dimer, an outcome supported by SAXS models. In contrast, Rac1 binds to monomeric HR1a, suggesting that this GTPase activates PKN1 via a different mechanism. These data provide structural insight into interactions between HR1 domains and the Rho family proteins and their potential consequences for PKN1 activation.

## Introduction

The Ras superfamily of small GTPases are often described as “molecular switches”, (reviewed in 1), which depend on regulator proteins including nucleotide exchange factors, GTPase activating proteins and guanine nucleotide dissociation inhibitors (reviewed in 2). Around 70 structures of GTPase-effector complexes have been solved, and the largest group of effectors utilise a pair of α-helices to interact with the G protein (reviewed in 3). The Rho family comprises 20 members, of which RhoA, Rac1 and Cdc42 are the best characterised. The filamentous proteins regulated by these three archetypal Rho GTPases are essential for cell migration (reviewed in 4; 5), for regulation of the cell cycle (6) and for control of vesicular trafficking (reviewed 7). RhoA, Rac1 and Cdc42 effectors involved in cytoskeletal regulation include the protein kinase C-related kinase (PKN) family (8), formins, ROCK proteins, TOCA, IQGAPs and N-WASP (9; 10; 11).

PKN1, PKN2 and PKN3 are members of the AGC serine/threonine kinase family, with a C-terminal catalytic domain (reviewed in 12; 13) and three N-terminal HR1 domains, HR1a, HR1b and HR1c (Figure 1A). These antiparallel coiled coils are effector binding domains for the Rho family GTPases, with the RhoA interaction with PKN1, via HR1a, activating the kinase (14; 15). This is thought to be due to release of autoinhibition by a pseudosubstrate sequence (16). Rac1 and RhoC have also been linked to increased PKN activity (17; 18). PKNs are also activated by phosphorylation by PDK1 (19; 20; 21), proteolysis by caspases and lipid binding (22; 8). PKNs contain a C2-like domain, which can facilitate plasma membrane recruitment (reviewed in 23), presumably allowing binding of lipids and Rho proteins. In PKN2, a ‘PKL’ region between the C2-like domain and the catalytic domain also mediates dimerization by interacting with the kinase domain, inhibiting the enzyme in *trans* (24). This region is conserved in PKN1 and PKN3, suggesting a similar inhibitory mechanism in all the PKNs.

**Figure 1.**
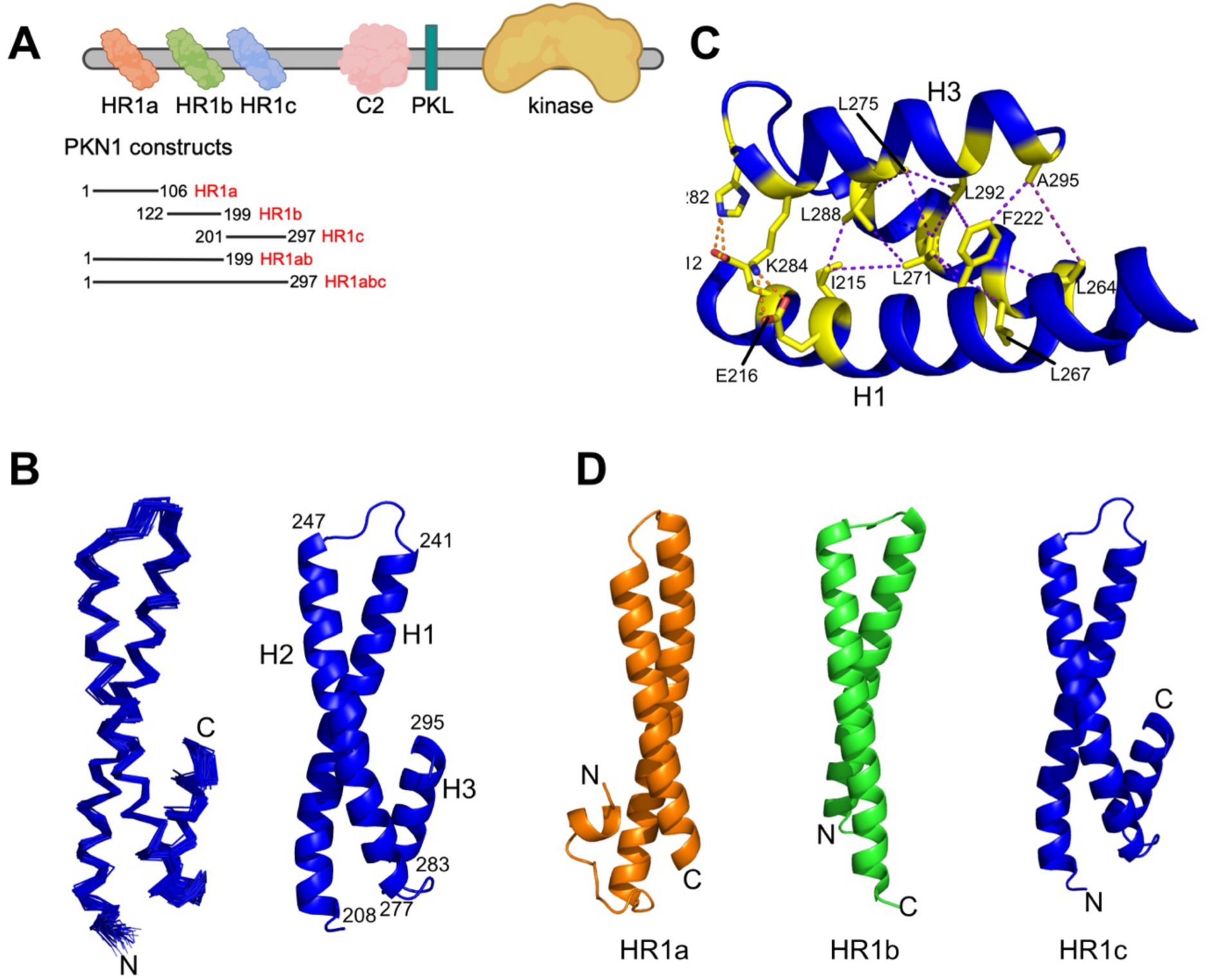
Solution structure of the PKN1 HR1c domain. (A) Domain structure of PKN1. The constructs used in this study are shown. (B) Structure of the PKN1 HR1c domain. Left: the ensemble of the 45 lowest energy structures of PKN1 HR1c (residues 206-297). Right: cartoon representation of the structure closest to the mean. The residue numbers indicate the boundaries of each helix. Helices are numbered H1-3. (C) Triple helical region of HR1c (coloured in blue). Residues involved in NOEs are shown as yellow sticks, with the atoms involved connected by dotted purple lines. The polar interactions between H1 and H3 are shown as dotted orange lines. (D) Structural comparison of PKN1 HR1 domains. HR1a (PDB:1CXZ) is coloured orange, HR1b green (PDB:1URF) and HR1c blue. N- and C-termini are labelled.

A role for PKNs has been described in cytoskeletal regulation (19; 25; 26; 18; 27) and cell adhesion (10). PKN1 and PKN2 have been implicated in androgen receptor signalling (28), and histone H3 Thr11 phosphorylation (29). PKN1 and PKN2 are important for mitotic entry and exit (30). PKN1 has also been implicated in the downregulation of the pyrin inflammasome response downstream of RhoA (31), whereas PKN2 has a distinctive role in development (32; 33) and PKN3 is involved in angiogenesis (34).

RhoA, RhoB and RhoC interact with PKN1, PKN2 and PKN3, indicating that the PKNs are effectors of the entire Rho subfamily (35; 18). The structure of RhoA in complex with PKN1 HR1a (36) identified two potential RhoA contact sites (I and II) for HR1a but mutational analysis defined contact site II as the relevant binding position (37). Rac1 binds with a high affinity to both HR1a and HR1b of PKN1 (38) and HR1b interacts with Rac1 at a site equivalent to the contact II site of RhoA (39).

No interacting partners have been identified for the HR1c domain and there are no reported structures. Furthermore, there is no structural information on the entire HR1 region or full length PKN. Here we have solved the structure of the PKN1 HR1c domain and established that some of the HR1 domains behave as oligomerization motifs. We have generated low resolution structural information on dimerized PKN1 domains and their complexes with the small GTPases RhoA and Rac1 and have investigated intra- and inter-molecular interactions between the three HR1 domains. The characterisation of these complexes supports a role for HR1-based dimerization in the regulation of PKN.

## Results

### The structure of the HR1c domain

The PKN1 HR1c construct used in this study comprised residues 201-297 (Figure 1A), which included a coiled coil domain and a putative extra α-helix, predicted by JPred in all three PKN isoforms (40). HR1c binding to the three archetypal Rho GTPases, RhoA, Rac1 and Cdc42 was tested but no interaction was observed (data not shown).

NMR experiments were performed and resonances assigned as described (41). A total of 2819 NOE restraints, derived from three-dimensional ^15^N-separated NOESY and ^13^C-separated NOESY experiments, were translated by ARIA into 2634 unambiguous restraints for structure calculations. After eight iterations there were 2026 unambiguous and 518 ambiguous NOEs. The lowest energy structures had no violations and were well-converged (Figure 1B, Table 1) except for the N-terminal 12 residues. These 12 residues, seven of which are cloning artefacts, are not shown in any structural figures.

PKN1 HR1c comprises three α-helices, connected by short loops. The first two helices interact to form an antiparallel coiled coil, resembling the structures of the other PKN1 HR1 domains. Helix 1 spans 34 residues (Leu208^H1^-Ala241^H1^), helix 2 spans 31 residues (Arg247^H2^-Glu277^H2^), while helix 3 at the C-terminus of the domain spans 13 residues (Pro283^H3^-Ala295^H3^). Helix 2 is straight whereas helix 1 wraps around helix 2 with a slight left-handed twist, akin to the other HR1 domains from PKN1 (36; 38), TOCA1 and CIP4 (42; 43). Helix 3 packs against the other two helices to form a short section of triple coiled coil.

The triple coiled coil is stabilized by interactions between all three helices (Figure 1C). The most extensive interactions involve the aromatic ring of Phe222^H1^, which is buried between Leu264^H2^, Leu267^H2^, Leu271^H2^, Leu292^H3^ and Ala295^H3^. The twist of helix 1 appears to favour the docking of the Phe222 ring into this hydrophobic pocket, positioning helix 3 alongside helices 1 and 2. Helices 2 and 3 form a network of hydrophobic interactions, for example, between Leu292^H3^, Leu271^H2^ and Leu275^H2^. In addition, there are hydrophobic contacts between Ile215^H1^, Leu271^H2^ and Leu288^H3^, which promote formation of the triple coiled coil structure. At the edge of the triple coiled coil there are polar interactions between Glu212^H1^-His282^H3^ and Glu216^H1^-Lys284^H3^, which are likely to fix the orientation of helix 3 with respect to the other helices. Despite the number of contacts, helix 3 has an average intensity ratio around 0.5 in heteronuclear NOE experiments, compared with helices 1 and 2, whose intensity ratios were around 0.7 (Supplementary Figure 1A), consistent with relatively well-ordered regions of the protein backbone. Analysis of the T_1_/T_2_ ratios for the three helices indicates that this is likely due to the orientation of helix 3 with respect to the overall diffusion tensor (Supplementary Figure 1B).

The relative lengths of the two helices in the coiled-coil vary between the different HR1 domains: in HR1a and HR1c, helix 1 is longer, while in HR1b helix 2 is longer (Figure 1D). In HR1a, a short N-terminal helix of just 5 residues contacts both of the main helices. The extra helix in HR1c is longer but also packs against both main helices (Figure 1D). Furthermore, the C-terminal HR1c helix is only slightly tilted with respect to the main two helices and forms a section of triple coiled coil, while the N-terminal HR1a helix is almost perpendicular to the coiled coil.

### HR1 domains as drivers of oligomerization

To understand the interactions between HR1 domains, we examined the NMR spectra of the PKN1 HR1ab didomain. We have previously shown that, like HR1c, HR1b is monomeric and well-behaved in solution (38). In contrast, the HR1ab spectra were of very poor quality, with just a few sharp peaks, which improved dramatically as the sample concentration was reduced stepwise from 360 μM to 15 μM (Supplementary Figure 2). This suggested that at higher concentrations the HR1ab protein was forming an oligomer that interconverted with the monomer, whereas at the lowest concentration the monomer was the predominant species. We therefore employed sedimentation velocity analytical ultracentrifugation (SV-AUC) to characterise the oligomerization of each PKN1 isolated HR1 domain, the HR1ab “didomain” and the HR1abc “tridomain”.

#### Oligomerization of PKN1

The sedimentation data for the PKN1 individual HR1 domains indicate that both HR1b and HR1c exist as a single species (Figure 2A), with molecular weights close to those expected (Supplementary Table 1A). In contrast, PKN1 HR1a exists as two species and the data fit best to a monomer-dimer model. Their masses cannot be accurately calculated because the sedimentation coefficients (s-values) of the two species vary with concentration, presumably due to a dynamic equilibrium between monomer and dimer that is concentration dependent. The proportions of dimer and monomer at the different concentrations (Supplementary Table 1A) indicate that dimerization is weak, with a *K*_d_ between 100 and 200 μM.

**Figure 2.**
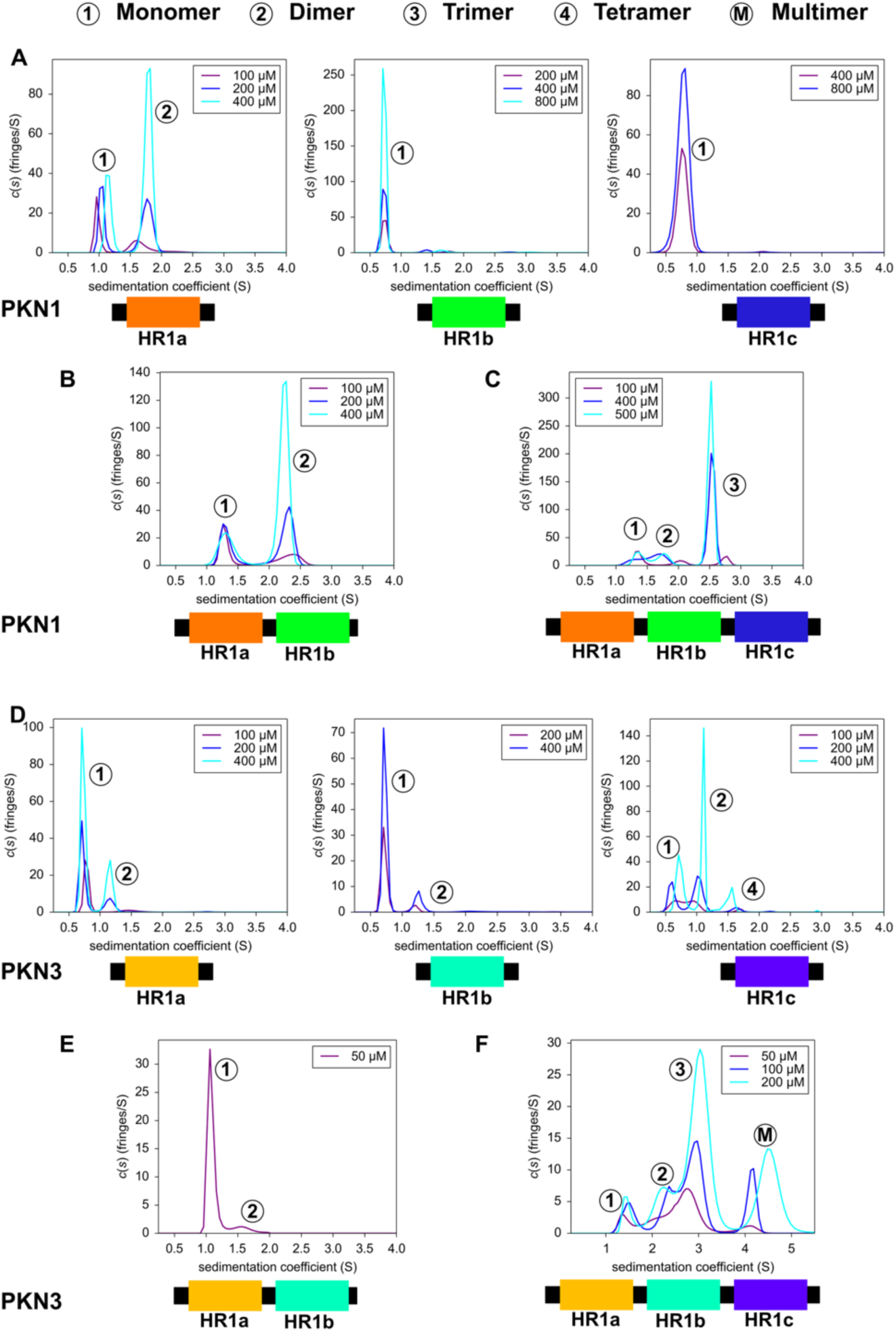
Oligomerization of HR1 domains measured by analytical ultracentrifugation. SV-AUC experiments were performed at the concentrations indicated and plotted as sedimentation coefficient distribution, c(s), *vs* the sedimentation coefficient, s. The numbers in circles next to each peak indicate the number of chains in that species based on the calculated molecular weights. M denotes a multimer whose size could not be accurately determined. Sedimentation profiles of (A) PKN1 HR1a, PKN1 HR1b and PKN1 HR1c single domains; (B) PKN1 HR1ab didomain; (C) PKN1 HR1abc tridomain; (D) PKN3 HR1a, PKN3 HR1b, PKN3 HR1c single domains. For HR1b, 100 μM data were excluded due to poor signal-to-noise; (E) PKN3 HR1ab didomain. This was unstable at concentrations higher than 50 μM; (F) PKN3 HR1abc tridomain.

The HR1ab didomain of PKN1 also dimerizes, with a similar *K*_d_ to that of HR1a alone (Figure 2B, Supplementary Table 1A) so that the presence of HR1b does not affect HR1a dimerization. The peak positions vary with concentration, consistent with monomer-dimer exchange, with smaller shifts than those of free HR1a, suggesting that exchange occurs on a slower timescale. Exchange between the monomer and dimer is in agreement with the NMR spectra recorded on the same construct (Supplementary Figure 2): at higher concentrations there is exchange at a timescale that renders the NMR peaks invisible, whereas at lower concentrations the monomer dominates and the spectra drastically improve.

Surprisingly, the HR1abc tridomain contained three exchanging species of molecular mass broadly corresponding to monomer, dimer and trimer, the trimer being the predominant species at higher concentrations (Figure 2C). As HR1ab does not form a trimer, this suggests that there are additional intermolecular/intramolecular interactions involving HR1c, for example that HR1c interacts with HR1a and/or HR1b.

In summary, SV-AUC indicates that HR1a and HR1c from PKN1 can both participate in interdomain interactions, facilitating dimerization in HR1ab and trimerization in HR1abc.

#### Oligomerization of PKN3

To understand whether the HR1 domains from other PKN proteins were also involved in multimerization, we investigated similar constructs from PKN3. SV-AUC experiments recorded show that all the individual PKN3 HR1 domains have some propensity to oligomerize (Figure 2D, Supplementary Table 1B). PKN3 HR1a and HR1b form weak dimers, with *K*_d_ values well above 400 μM. PKN3 HR1c also dimerizes but a small amount of tetramer was also observable. At the lowest concentration where the tetramer is barely populated, the monomer:dimer ratio is close to 1:1, suggesting a similar *K*_d_ to that of PKN1 HR1a. This suggests that in PKN3 the HR1c domain is the driver of oligomerization. The concentration dependent s-values indicate that PKN3 HR1c is in a dynamic equilibrium between different oligomeric states, in agreement with the broad signals in NMR experiments (data not shown). PKN3 HR1c has a similar helical content as the PKN1 HR1c, based on CD analysis (Supplementary Figure 3A) but the two domains have very different thermal stabilities (Supplementary Figure 3B). PKN1 HR1c shows a single cooperative transition in its melting curve, consistent with a small monomeric domain, whereas the PKN3 HR1c has a shallower melting curve, suggesting the presence of multiple species that unfold at different rates/temperatures.

PKN3 HR1ab formed some dimer even at 50 μM (Figure 2E). This protein was unstable at higher concentrations but when HR1c was present (as HR1abc), the protein could be concentrated to 200 μM (Figure 2F). In the HR1abc tridomain, 4 species could be observed, with molecular mass estimation and s-value variation suggesting that HR1abc exists in a linked equilibrium of monomer, dimer, trimer and a larger oligomer that fits best to a pentamer (Figure 2F). The monomer was present at less than 15% even at the lowest concentration, indicating a higher propensity to multimerize but *K*_d_ values could not be estimated in this complex mixture.

### Interactions between PKN1 HR1c and other HR1 domains

The additional oligomeric states formed by both tridomains, compared with the HR1 single and HR1ab double domains, suggest that there are additional interdomain interactions involving HR1c. As the AUC data suggested that there are intermolecular or intramolecular interactions between PKN1 HR1c and HR1a/HR1b, we searched for contacts between isolated domains. In HR1abc the domains would be tethered by HR1a-mediated dimerization, which brings the other HR1 domains from two monomers into close proximity, bolstering low affinities. We therefore used NMR chemical shift perturbation, which can map even very weak binding between the domains.

Unlabelled PKN1 HR1a was titrated into ^15^N-labelled PKN1 HR1c and an ^15^N HSQC experiment recorded at each titration point (Figure 3A). Small chemical shift changes were observed for some HR1c backbone amide resonances upon HR1a titration, which was expected for a weak interaction involving a helical protein, where the backbone amides are buried. All the peaks were in a fast exchange regime and the titration endpoint was not achieved at a 1:1 ratio, implying that at the concentrations used (300 μM), there was still unbound HR1c present. Chemical shift changes that were larger than the mean + 1 SD map to residues on one face of helix α1 and in the short loop between helices 1 and 2 on the HR1c structure (Figure 3B and C(i)).

**Figure 3.**
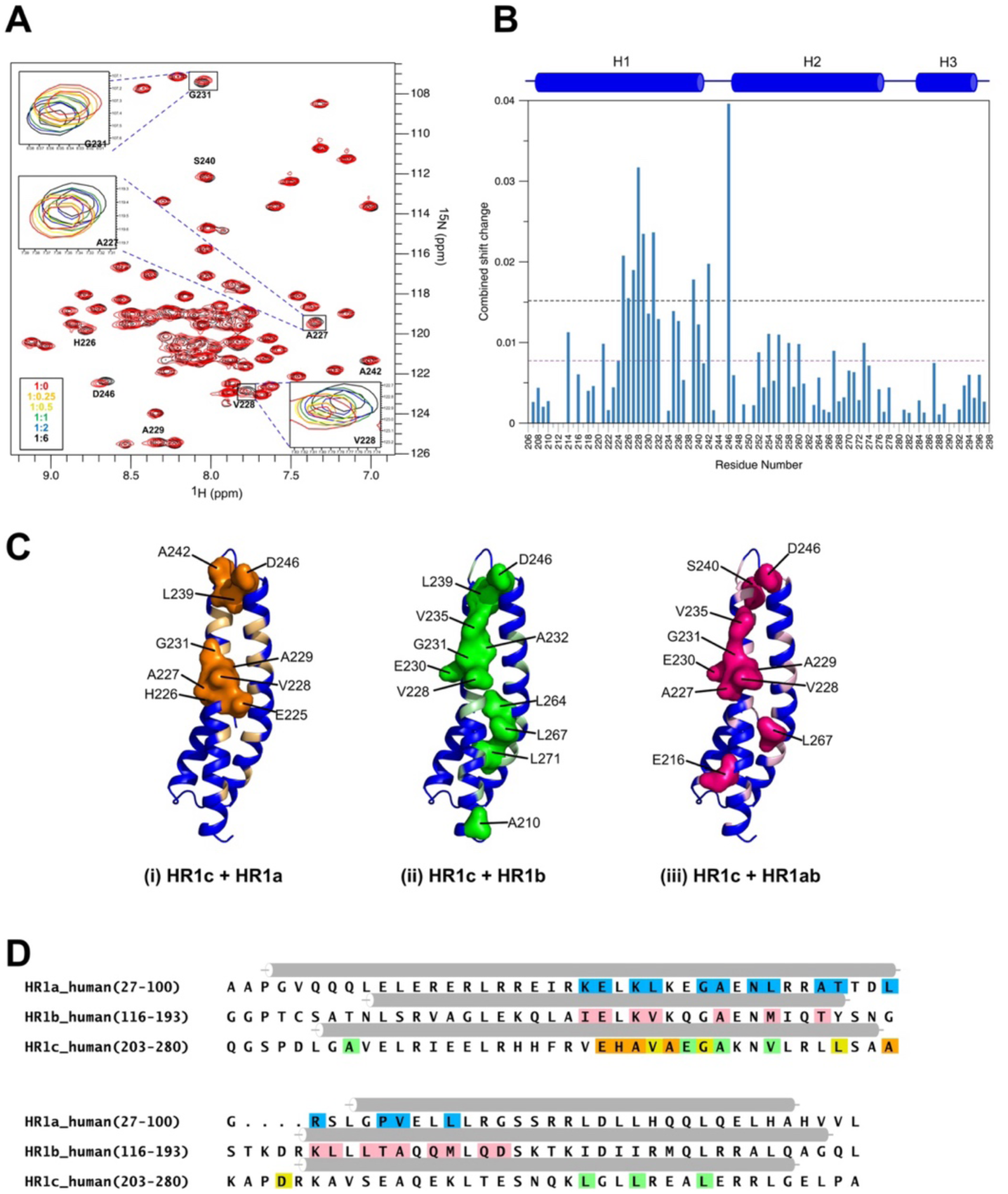
HR1c interactions with HR1a and HR1b. (A) ^15^N HSQC spectrum of ^15^N-labelled PKN1 HR1c free (red) and in the presence of PKN1 HR1a (black). Isolated resonances that shifted are labelled with their assignment and examples shown in the magnified insets, with the molar ratios 1:0 (red), 1:0.25 (orange), 1:0.5 (yellow), 1:1 (green), 1:2 (blue) and 1:6 (black). (B) Combined chemical shift changes at a 1:6 molar ratio. The average and average + one standard deviation are marked with purple and black dotted lines respectively. (C) HR1c residues whose resonances shifted by greater than average + one standard deviation are shown in a surface representation on the structure, when titrated with (i) HR1a (orange), (ii) HR1b (green) and (iii) HR1ab (magenta). Residues whose resonances shifted only above the mean are coloured on the HR1c structure in light orange, light green and pink for HR1a, HR1b and HR1ab titrations, respectively. (D) Sequence alignment of PKN1 HR1 domains. Interacting residues in the RhoA-HR1a (Maesaki *et al.*, 1999) and Rac1-HR1b (Modha *et al.*, 2008) complexes are coloured blue and red, respectively. HR1c residues classed as HR1a-interacting are coloured orange, those that interact with HR1b only are in green, those that interact with both HR1a and HR1b are coloured yellow.

To investigate whether HR1b also interacts with HR1c, unlabelled HR1b was added to ^15^N-labelled HR1c and HSQC experiments recorded. Even smaller chemical shift changes were observed (Supplementary Figure 4A), with significant chemical shift changes (greater than average + 1 SD) between residues 228 and 246, overlapping the HR1a binding site (Figure 3C(ii)). A second region, within helix 2, also exhibited significant chemical shift changes, indicating that the residues perturbed by binding HR1a and HR1b are not identical (compare Figure 3C panels (i) and (ii)). Titrations of the HR1ab didomain with HR1c showed similar shift changes to those with HR1b alone (Supplementary Figure 4, Figure 3C(iii)). This implies that HR1c preferentially interacts with HR1b in the context of the didomain, although given the similarity of the residues whose shifts change, it cannot be excluded that HR1a and HR1b compete for binding to HR1c.

When the chemical shift changes in HR1c are compared to the residues in HR1a and HR1b that interact with RhoA and Rac1 respectively, it is clear that they partially overlap (Figure 3D). This suggests that if a GTPase were to bind to HR1c, its binding would be competitive with that of HR1a/HR1b, which would lead to conformational changes in the HR1abc region.

### Structural insight into dimerization of PKN1 HR1 domains

The equilibria between monomeric and dimeric forms presented a challenge for determining high-resolution structures of PKN1 dimers, so to obtain useful structural information, SEC-SAXS (size exclusion chromatography small angle X-ray scattering) experiments were recorded on PKN1 HR1a, HR1ab and HR1abc. SAXS data for the dimeric forms of all three proteins were obtained but the HR1abc trimer was not sufficiently populated to be detected at the concentrations available. *Ab initio* shape reconstruction resulted in models whose SAXS patterns are in excellent agreement with the experimental SAXS patterns. (Supplementary Table 2).

#### PKN1 HR1a

The HR1a dimer is an elongated structure with a maximum diameter of 97 Å (Figure 4A; Supplementary Figure 5A). The higher density of the SAXS envelope at the ends suggests that the N- and C-terminal extensions are located there, and that the dimerization interface involves the inter-helical loop and the ends of the helices nearest to the hairpin loop. Using the HR1a structure from the RhoA-HR1a complex (PDB:1CXZ), we performed SAXS-guided docking of the HR1a dimer using FOXSDOCK, followed by MULTIFOXS to allow the flexible termini to move (44; 45). The resulting model had a χ^2^ of 1.05 (Supplementary Table 2). We also predicted the structure of the HR1a dimer using the AlphaFold3 server (46). The top model has an ipTM (interface predicted template modelling) score of 0.66, indicating that it may be correct and MULTIFOXS (to allow for flexibility in the termini) yielded a χ^2^ of 1.01. The two models were similar, with a backbone RMSD of 2.2 Å, and were of an offset head-to-head dimer, with a shape that matched the SAXS envelope (Figure 4A). The *R*_g_ of the models were 28-30 Å, slightly lower than the ∼32 Å obtained from Guinier and p(r) analysis. This may indicate that the short N-terminal helix of the HR1a domain, which was treated as rigid, has some flexibility. Overall, the low χ^2^ of the fit to the SAXS data, and their agreement with both the shape of the SAXS envelope and each other indicate that they are representative of the HR1a dimer in solution.

**Figure 4.**
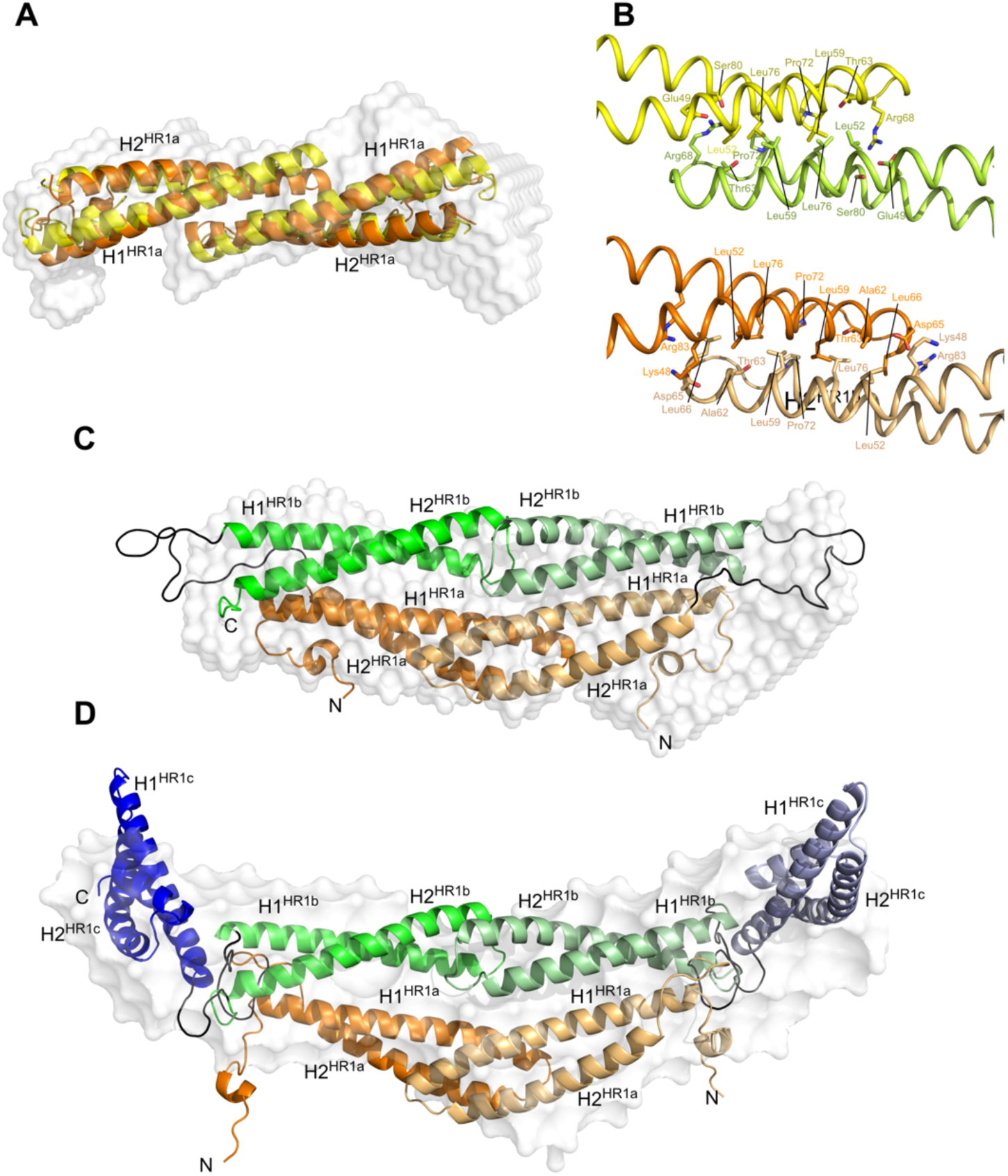
Insights into HR1a-mediated dimerization of PKN1. (A) HR1a dimer predicted by AlphaFold3 (yellow) or FOXSDOCK (orange) overlaid over the *ab initio* SAXS model (white). (B) Close-up of sidechain interactions between the HR1a domains from AlphaFold3 (yellow and green) and FOXSDOCK (shades of orange). (C) HR1ab HADDOCK model overlaid over the *ab initio* SAXS model (white). (D) HR1abc HADDOCK model overlaid over the *ab initio* SAXS model (white). In panels (C) and (D), HR1a is shown in shades of orange, HR1b in green and HR1c in blue. The linkers between the domains are black. One monomer is shown in dark colours and the other in pale shades. The N-terminal helix of each domain is labelled H1 and the C-terminal helix labelled H2, with the HR1 domain identified by the superscript.

Although the details of the two HR1a dimer models differ, the interface between monomers is broadly similar, with a hydrophobic core comprising residues Leu52^H1^, Leu59^H1^, Pro72^H2^ and Leu76^H2^ (Figure 4B). This core is flanked by polar residues that form salt bridges/hydrogen bonds that vary in the models: either Lys48^H1^/Arg83^H2^ to Asp65^H2^ or Glu49^H1^/Ser80^H2^ to Arg68^linker^.

#### PKN1 HR1ab

The SAXS envelope of the HR1ab didomain indicates that this protein also forms an elongated shape (Figure 4C; Supplementary Figure 5B). AUC showed that the dimerization of HR1ab is driven by HR1a association (Figure 2A and B), hence the core of the structure is likely to be formed by an HR1a dimer with the HR1b domains located on the exterior. As the maximum diameter of the HR1ab dimer envelope is only 20 Å bigger than HR1a (Supplementary Table 2), the HR1b domains are not fully extended away from the HR1a dimer, despite the 20-residue linker between the domains. This suggests that there are intramolecular interactions between HR1a and HR1b within each monomer.

To construct a model of the HR1ab dimer the chemical shift differences between free HR1b and HR1b in the context of the HR1ab didomain (Supplementary Figure 6) were used to provide restraints for HADDOCK2.4 (47). Interactions between the HR1a and HR1b domains were assumed to be intramolecular, as the chemical shifts of HR1ab came from a sample that was diluted to 15 μM and was therefore monomeric. There were 15 clusters resulting from the HADDOCK calculations, and the cluster with the best HADDOCK score also fit the SAXS data best, with an average χ^2^ of 3.1 across 23 models and a minimum of 2.2. After remodelling the linker between the domains, the final model had a χ^2^ of 1.51, *R*_g_ of 33.4 Å and *D*_max_ around 120 Å (Supplementary Table 2), which are similar to the values calculated from the SAXS data. The resulting model also fits the SAXS *ab initio* envelope reasonably well (Figure 4C). The model shows extensive contacts between HR1b and HR1a within a single chain, with the domains lying parallel to each other and forming a four-helix bundle. The two HR1b domains in the dimer are relatively close to each other but do not form significant contacts.

The HR1ab dimer was also predicted by AlphaFold3 but the ipTM score was 0.36, which implies that the prediction may have failed. All five models predicted by the AF3 server were tested against the SAXS data using FOXS. Four of the five models were similar and had χ^2^ values of 4.7-5.2 whereas one model had a distinct domain orientation and a χ^2^ of 1.96. This model was of a similar overall shape to the HADDOCK model, but the orientation of HR1b relative to HR1a was different, so that fewer HR1b residues whose chemical shifts had changed in the presence of HR1a were in contact with HR1a (Supplementary Figure 6C).

### PKN1 HR1abc

Dimerization of the HR1abc tridomain is presumably also mediated by self-association of the HR1a domain. SAXS analysis of the HR1abc dimer yielded a maximum diameter of 163 Å, indicating an extended conformation (Supplementary Table 2). Assuming that the HR1a dimer is at the core of the structure, HR1c has extended the length of the HR1ab dimer by ∼46 Å. Each HR1c monomer is ∼55 Å, so these domains are likely to be partially extended from the core, with the HR1b domains forming a bridge between HR1a and HR1c.

The HR1abc dimer was built from the HR1ab dimer model and our HR1c structure, using the NMR titration of HR1ab into HR1c (Supplementary Figure 4B) to provide restraints. There were 10 clusters resulting from the HADDOCK calculations, and within the cluster with the best HADDOCK score was the model that fit the SAXS data best, with a χ^2^ of 3.67. After refining the N-terminal extension and the loops between HR1 domains using Modeller, the final model fit the SAXS data with a χ^2^ of 1.67. This model fits the SAXS *ab initio* envelope reasonably well (Figure 4D), although HR1c extends beyond the envelope. The SAXS envelope represents the average structure in solution, whereas the single model from HADDOCK resembles one conformation. This implies that the position of HR1c with respect to the HR1ab domains is not fixed and there is some flexibility in their interactions. The model does not fit all the residues identified by the NMR titration data, but neither did any of the HADDOCK models. This is either because chemical shifts are very sensitive to subtle reorientations and therefore often over-estimate binding interfaces or is another indication that there is flexibility in the interaction between HR1c and HR1ab. Compared with the interactions between HR1a and HR1b, which lie parallel to each other and form close contacts, HR1c is at the edge of the HR1ab four-helix bundle and its coiled-coil approximately perpendicular to the HR1a and HR1b coiled-coils. Most of the contacts are between HR1b and HR1c, which is consistent with the chemical shift mapping data (Supplementary Figure 4), which shows that HR1ab titrations into HR1c look most similar to the equivalent HR1b titrations.

The HR1abc dimer was also predicted by AlphaFold as before, but the five resulting models were all in different orientations and have ipTM scores around 0.2, indicating that the prediction has failed. The Alphafold models did not fit the SAXS data well, with χ^2^ values of 15-16.

### Binding of Rho Proteins to HR1 dimers

As RhoA binds to HR1a and activates PKN1, it was of interest to investigate whether RhoA affects oligomerization. Comparison of the HR1a dimer model with the structure of the complex formed between RhoA and HR1a shows that the binding site for RhoA overlaps with the dimer interface, so that Rho binding is expected to break the HR1a dimer (Figure 5A).

**Figure 5.**
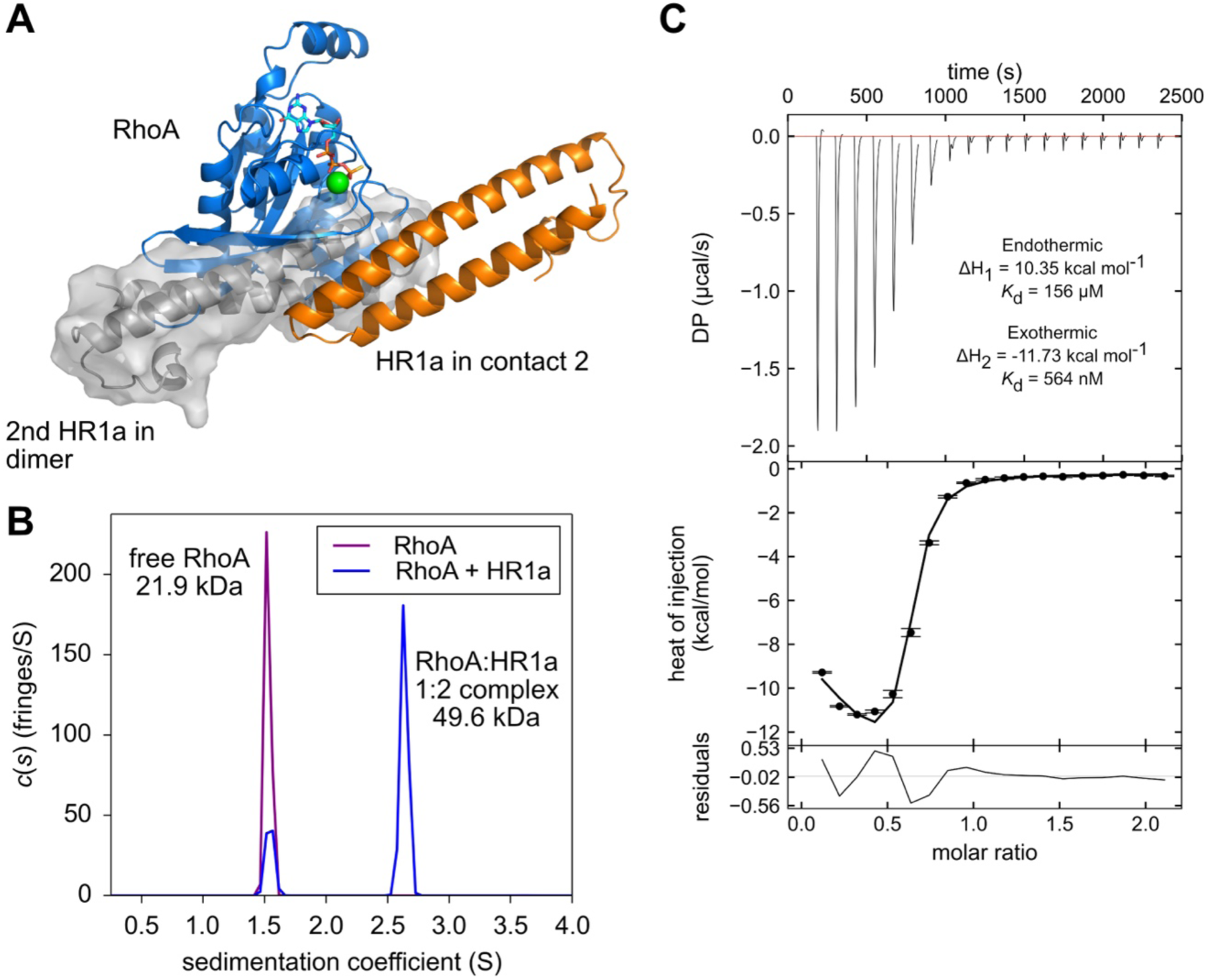
RhoA binds to dimeric HR1a. (A) The free HR1a dimer and the RhoA/HR1a complex are mutually exclusive as shown by the structural clash. RhoA is shown in blue, the Rho-interacting HR1a domain is shown in orange and the second HR1a domain in the HR1a dimer is shown in grey. (B) Sedimentation profile of free RhoA (150 μM) and RhoA mixed with PKN1 HR1a in a 1:1 ratio (150 μM of each protein). RhoA forms a 1:2 complex with PKN1 HR1a. (C) The binding of RhoA to HR1a measured by isothermal titration calorimetry. Heat changes were recorded as 1 mM RhoA was titrated into 0.1 mM HR1a (monomer concentration). Two events were observed, which were fit simultaneously to give an overall χ^2^ of 1.03.

AUC experiments were performed with (a) free RhoA·GMPPNP and (b) an equimolar mixture of RhoA·GMPPNP and HR1a. Both proteins were at 150 μM, close to the dimerization *K*_d_ of HR1a, so that the amount of HR1a monomer and dimer available would be near equal (Figure 2A; Supplementary Table 1A). Data for free RhoA fit to a molecular weight of 21.9 kDa (Figure 5B), close to the expected mass of 21.7 kDa. In the equimolar mixture two species were detected: one has the same s-value as free RhoA whereas the second fits to a molecular weight of 49.6 kDa, consistent with a 1:2 complex of RhoA:HR1a (molecular weight 46.4 kDa). This is supported by the observation of free RhoA in the mixture, despite the equimolar ratio used in the experiment. This complex therefore comprises either RhoA bound to a reoriented HR1a dimer or RhoA bound to two HR1a monomers. No free HR1a monomer was observed in the sedimentation profile (Figure 2A, Supplementary Table 1A), indicating that all detectable HR1a was in complex with RhoA.

The interaction between RhoA and HR1a was also assessed by isothermal titration calorimetry (ITC). RhoA was injected into the ITC cell containing 100 μM PKN1 HR1a, which at this concentration is expected to be partially dimeric. The data showed that two separate events occurred upon titration (Figure 5C). One was endothermic, with a *K*_d_ of 156 μM, whereas the second event was exothermic with a much lower *K*_d_ of 564 nM. The *K*_d_ of the second event is similar to that of HR1a-RhoA measured by other methods (38; 48). An ITC experiment performed with HR1a at 15 μM, well below the dimer *K*_d_, gave the same, albeit noisier, results (data not shown), arguing against dimer dissociation being responsible for the endothermic process. The stoichiometry of the interaction could not be accurately determined due to the complex set of events, but it is clearly not 1:1, with the mid-point of the isotherm being around 0.5.

To understand how RhoA binds to two HR1a molecules the complex was characterised by SEC-SAXS (Figure 6A; Supplementary Figure 7; Supplementary Table 2). The SEC profile of an equimolar sample of the two proteins indicated that there were two species present (Supplementary Figure 7A). The molecular mass calculated from the SAXS data corresponding to the second peak was ∼22 kDa (Supplementary Table 2) and the SAXS envelope matches the RhoA structure (Supplementary Figure 7D). The other species, with an estimated molecular mass of ∼55 kDa (Supplementary Figure 7B), was assigned to the RhoA:HR1a 1:2 complex. The *ab initio* SAXS model consists of two lobes in a V-shape, one of which fits RhoA, the other therefore containing the HR1a domains (Figure 6A).

**Figure 6.**
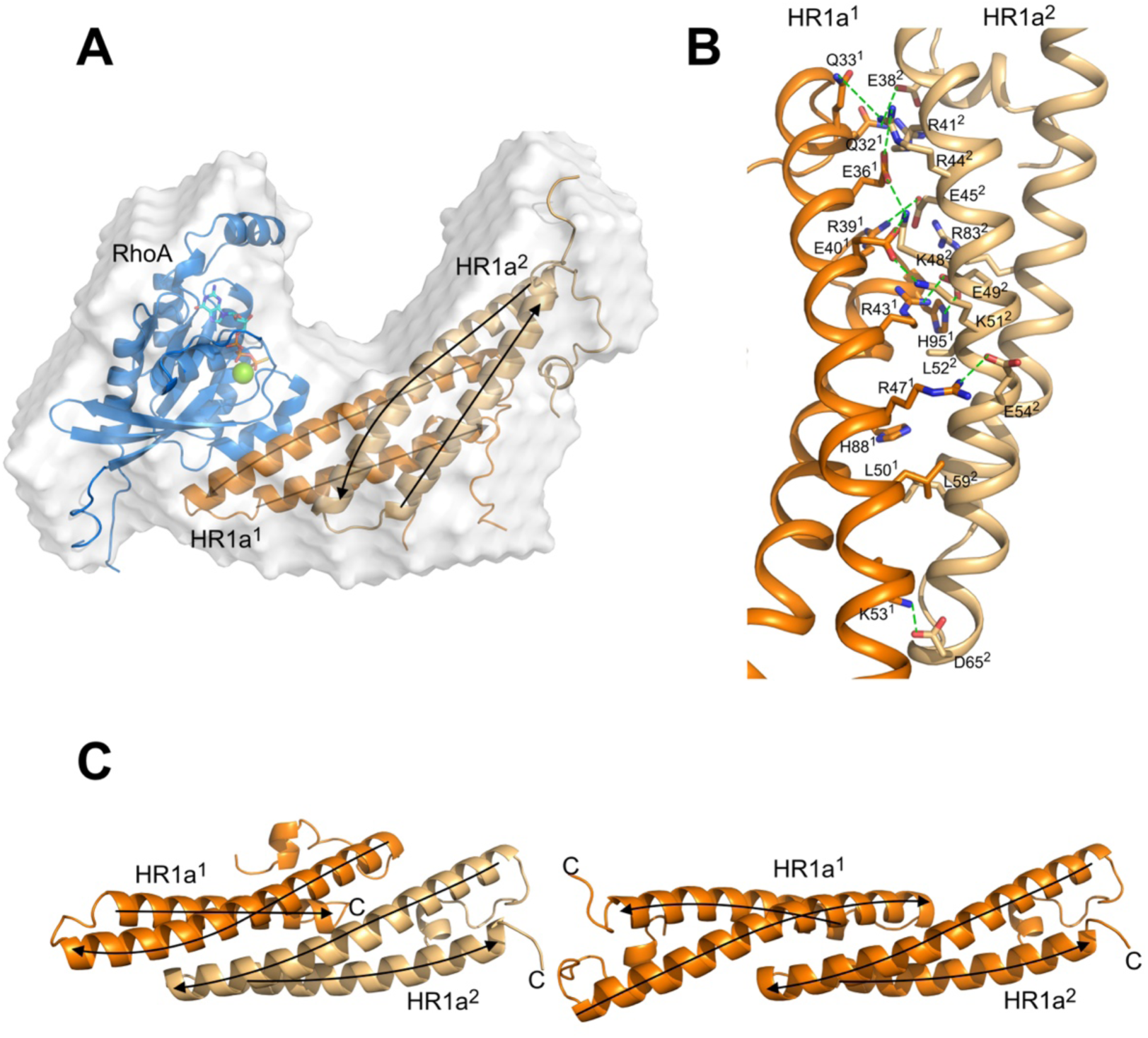
Structural insight into the 1:2 RhoA:PKN1 HR1a complex. (A) RhoA:HR1a complex from FOXSDOCK overlaid over the low-resolution *ab initio* model (white surface). RhoA is shown in blue, GTP is shown in a stick representation, Mg^2+^ is a green sphere, HR1a is shown in orange, with the monomer that interacts with RhoA in a darker shade. (B) Close-up of the interface between the two HR1a domains in the RhoA-bound HR1a dimer. HR1a^1^ contacts RhoA and is coloured dark orange. Interacting side chains are shown in a stick representation, with carbon orange, oxygen red and nitrogen blue. Predicted ionic interactions/hydrogen bonds are indicated by green dashed lines. (C) Comparison of HR1a dimer bound to RhoA (left) and free HR1a dimer (right). The models were overlaid on HR1a^2^. The RhoA-bound dimer is coloured as in panels A and B. Arrows indicate the chain direction (N- to C-) within each α-helix.

Assuming that one HR1a domain interacts with RhoA at the contact II site (36), the other HR1a domain must be interacting with either the bound HR1a or with RhoA. We therefore used FOXSDOCK for SAXS-guided docking of a free HR1a domain onto the RhoA-HR1a structure (PDB:1CXZ; contact II from PDBePISA). The best-fit model had the second HR1a domain (HR1a^2^) contacting the HR1a domain (HR1a^1^) already bound to RhoA (Figure 6A; Supplementary Table 2) and after refinement of the tails the χ^2^ was 2.23. The overall shape matches that of the SAXS envelope well and shows that the second HR1a domain (HR1a^2^) uses its RhoA-binding region to interact with HR1a^1^, which in turn is bound to RhoA (Figure 6A). This implies that RhoA binding by HR1a^1^ causes a subtle rearrangement that allows HR1a^2^ to bind at a newly created interface on HR1a^1^ (Figure 6B). The most obvious difference between the RhoA-bound and free HR1a dimers is that the HR1a domains in the free dimer are antiparallel and only interact along approximately half their length, whereas in the RhoA-bound dimer the two HR1a domains are parallel and interact along more of the length of the coiled-coils (Figure 6C).

In the RhoA complex, the interface between HR1a^1^ and HR1a^2^ buries ∼1900 Å^2^, in contrast to the free HR1a dimer model, where ∼1700 Å^2^ are buried. The free HR1a dimer interface is relatively hydrophobic with flanking polar contacts (Figure 4B), and involves both helices of the coiled-coil, whereas the RhoA-bound HR1a^1^-HR1a^2^ model interface is dominated by polar interactions and mostly comprises interactions between residues in the first helix (Figure 6B).

The data from AUC, ITC, SAXS (envelope, docking and modelling) experiments and published binding data, are all in agreement with the FOXSDOCK model. Alternative binding modes for RhoA binding to two HR1a domains were explored but these were incompatible with the SAXS data of the complex (Supplementary Figure 8). The best model from AlphaFold3 predicted that each HR1a monomer forms a single long helix, and that two of these come together to form a dimer that interacts with RhoA. This arrangement did not fit with the SAXS data and had an iPTM score of 0.55.

Analysis of SEC-SAXS experiments performed on RhoA-HR1ab and RhoA-HR1abc complexes resulted in *R*_g_ and MW values consistent with a 1:2 complex of RhoA with the didomain and tridomain (Supplementary Figure 9; Supplementary Table 2). A model of the RhoA-HR1ab complex was built from the RhoA-HR1a model and refined in MULTIFOXS to a single state model where the HR1b domains point away from the rest of the complex (Supplementary Figure 9B; Supplementary Table 2). This is consistent with the observation that HR1b does not contribute to the interaction with RhoA (37). The RhoA-HR1abc 1:2 complex is also extended based on the D_max_ value (Supplementary Table 2), suggesting that like HR1b, HR1c extends away from the centre of the structure. The calculated envelope density represents a population average, and the positions of the domains cannot be accurately defined (Supplementary Figure 9D). The two HR1c domains appear to be mobile and independent from the rest of the complex preventing any further models being built.

### Rac1 stabilises monomeric PKN1 HR1a

Rac1 interacts with both HR1a and HR1b (38) and the solution structure of the HR1b complex showed that they interacted with a 1:1 stoichiometry (39). SEC-SAXS data was recorded on Rac1 with HR1a and surprisingly this was present as a 1:1 complex, implying that Rac1 stabilises the HR1a monomer (Supplementary Figure 10A; Supplementary Table 2). A model of HR1a bound to Rac1 at the site equivalent to the HR1b binding site was generated using Modeller and refined against the SAXS data using MULTIFOXS. The resulting model fitted the SAXS envelope well (Figure 7A).

**Figure 7.**
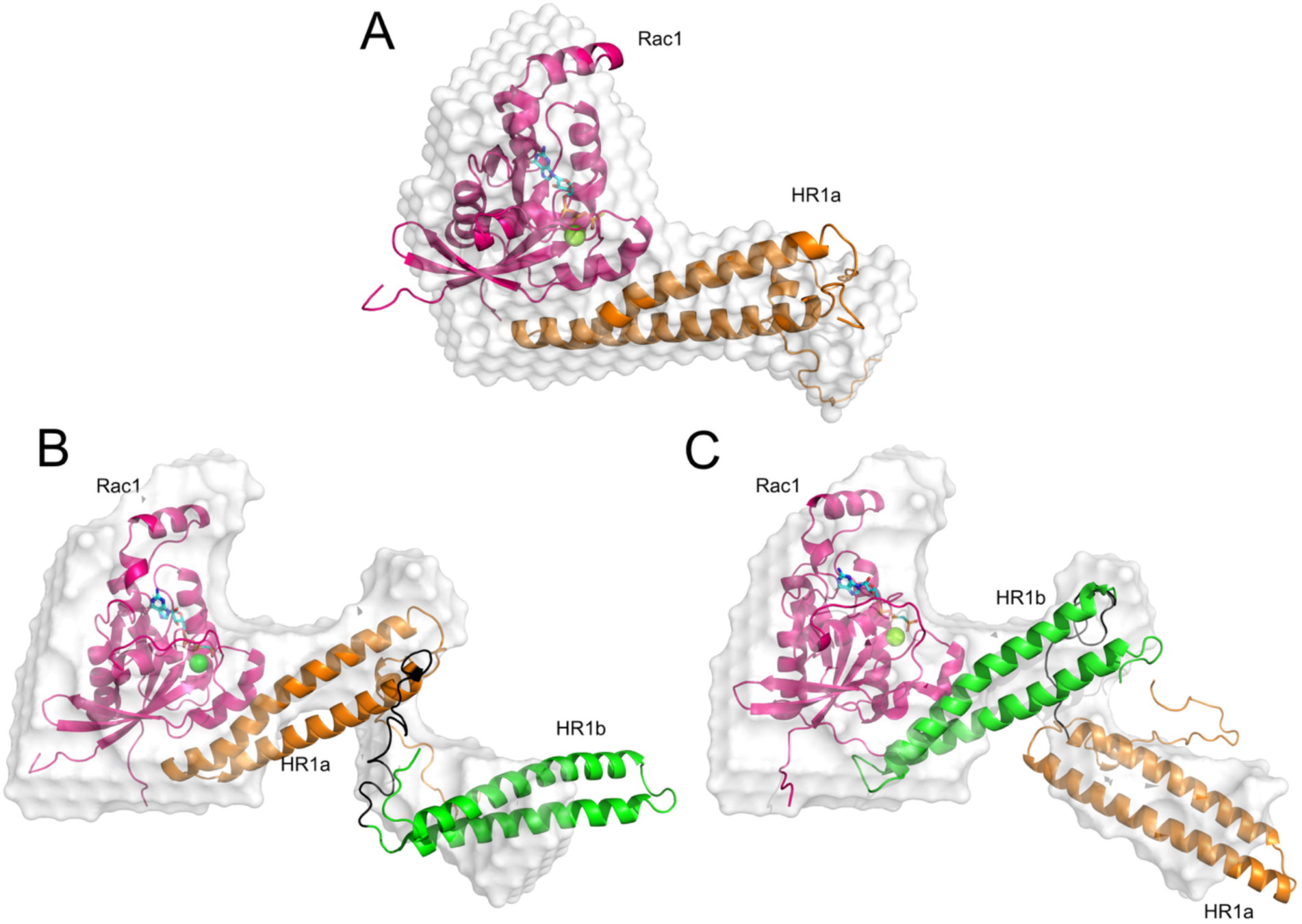
Structural insight into Rac1:PKN1 HR1 complexes. (A) Rac1:HR1a complex model overlaid over the low-resolution *ab initio* model (white surface). Rac1 is shown in pink, GTP is shown in a stick representation, Mg^2+^ is a green sphere, HR1a is shown in orange. (B,C) Rac1:HR1ab complex models when HR1a (B) or HR1b (C) contact Rac1. Models are shown overlaid over the low-resolution *ab initio* model (white surface). Cartoon colours as in panel A, with HR1b shown in green.

Rac1 binds to both HR1a and HR1b, with no evidence of cooperativity (38). To investigate whether Rac1 binding to HR1b allows a free HR1a to dimerize, or a single HR1ab didomain can bind two Rac1 molecules, Rac1 and HR1ab were mixed in a 2:1 ratio and characterised by SEC-SAXS. The SAXS analysis shows that this mixture also results in formation of a 1:1 complex (Supplementary Figure 10B), so that HR1a does not dimerize in this context, and only one Rac1 molecule binds per HR1ab dimer. The SAXS envelope shape implies that one HR1 domain interacts with Rac1, while the other points away from the G protein. Given the similar shape and affinities of the HR1 domains, it was not possible to determine whether one binds preferentially or the two domains compete for the same binding site on Rac1. Models with HR1a or HR1b interacting with Rac1 both fit very well to the scattering data and reasonably well to the low-resolution SAXS envelope (Supplementary Table 2; Figure 7B,C). However, the interaction of HR1a with Rac1 leads to dissociation of the free HR1a dimer and, since the monomer persists in the HR1ab-Rac1 complex, we speculate that HR1a is bound to Rac1.

## Discussion

We show here that PKN1 HR1c is an antiparallel coiled coil with a left-handed twist, consistent with the structures of other HR1 domains (36; 38; 43; 42) but with an additional third, short helix that packs against the other two helices (Figure 1B). Although no GTPase binding partner has yet been identified for PKN1 HR1c, there are some GTPase effectors whose coiled coil GTPase binding domains (GBD) contain three helices *e.g.* the mDia1 GBD consists of two helices that interact with the switch regions of RhoC in a GTP-dependent manner (49). A third helix packs against these helices and stabilises their orientation, promoting GTPase binding. The C-terminal helix of HR1c, which also packs against the two main helices, could have a similar role stabilising the coiled coil orientation in protein-protein interactions.

It is feasible however that the role of HR1c is not GTPase binding but is instead to promote the formation of PKN1 oligomers. In isolation, PKN1 HR1a and HR1ab can dimerise, in contrast to the monomeric HR1b and HR1c domains. Addition of HR1c to form the HR1abc tridomain results in formation of a trimer (Figure 2C), indicating that extra intramolecular and intermolecular interactions exist in the tridomain and that HR1c has a role in mediating these interactions. SAXS-based models, backed up by Alphafold predictions, showed that the HR1a domains arrange in a staggered anti-parallel dimer, and this dimer is likely to be at the core of the HR1ab and HR1abc proteins (Figure 4). We have also demonstrated weak interactions between pairs of isolated HR1 domains (Figure 3). As the HR1abc dimer is based on HR1a dimerization and if HR1abc oligomerization follows a linear pathway from monomer to dimer to trimer, it seems likely that the HR1abc dimer interacts with an additional protomer via HR1c. This trimer formation may however be a consequence of removal of HR1abc from the context of the full protein. AlphaFold3 confidently predicts an interaction between the C2-like domain and HR1c for all three PKNs (Supplementary Figure 11), suggesting that HR1c could be involved in contacting other regions of the protein.

Even at low concentrations PKN3 HR1abc formed a trimer and a higher order multimer, suggesting a higher degree of multimerization than PKN1. In PKN1, HR1a drives HR1ab dimerization and HR1c, although monomeric alone, supports trimerization when all three HR1 domains are present. In contrast, all PKN3 HR1 domains can dimerize alone but HR1c can also form tetramers and drives higher order oligomers in the context of the tridomain. It is therefore likely that there are subtle differences in the intermolecular and intramolecular interactions between the HR1 domains in the PKN1 and PKN3 oligomers. It is HR1a that drives PKN1 dimerization but in PKN3 the affinity of HR1a dimers is lower (compare Figure 2A and 2D) and the oligomerization is driven instead by HR1c. Given that RhoA binding to PKN1 HR1a reorients the HR1a dimer, the lower affinity of the PKN3 HR1a dimer means that a similar reorientation would be easier to achieve with PKN3. This, together with the different oligomerization states of PKN1 and PKN3 may provide a structural and mechanistic rationale for the non-redundant biological functions of these two PKNs, e.g. PKN3 but not PKN1 is required for bone resorption downstream of RhoA (51).

It is likely that interactions between the HR1 domains are important in regulating PKN catalytic activity, helping to maintain PKN1 in a ‘closed’, autoinhibited state. This is consistent with studies demonstrating that the N-terminal region of the PKNs hinders their catalytic activity, possibly by preventing recruitment of the activating kinase, PDK1 (8). Despite the relatively weak HR1a dimerization affinity, the *trans* domains would be tethered by intermolecular contacts between the PKL and kinase domain, which are in *trans* (24). Clearly there are multiple oligomerization sites in PKNs, which would explain the observation of higher order multimers when PKN1 was cross-linked (24). The AlphaFold3 prediction for PKN1 suggests an interaction between the PKL and kinase domain (Supplementary Figure 11), including basic residues in the PKL, which, when mutated increased the kinase activity (24). Although this interaction was not predicted as confidently as the HR1c-C2-like interaction, PKN2/3 AlphaFold3 models were similar. The basic residues in the PKL are predicted to form electrostatic contacts with acidic residues in the kinase domain near to the ATP binding pocket, suggesting a plausible mechanism for inhibition. The linker between the PKL and the kinase is long enough to allow the PKL-kinase interaction to occur in *trans*, which was suggested to be the case (24).

We used our HR1abc data-driven docked model in combination with the full-length AlphaFold prediction to generate a model of full-length PKN1 dimer (Figure 8). In the HR1abc dimer model, HR1c would be free to interact with the C2-like domain in the manner confidently predicted by AlphaFold (Supplementary Figure 11). The two models were overlaid on HR1c, so that the N-terminal ∼300 residues came from the HR1abc data-driven dimer model, and the remainder of the protein came from AlphaFold3. The model has the HR1a dimer in the centre with HR1c and the C2-like domain on the periphery, and the kinase active site shielded by the PKL region. The long linkers between the PKL and the kinase allow the PKL-kinase interaction to occur in *trans*.

**Figure 8.**
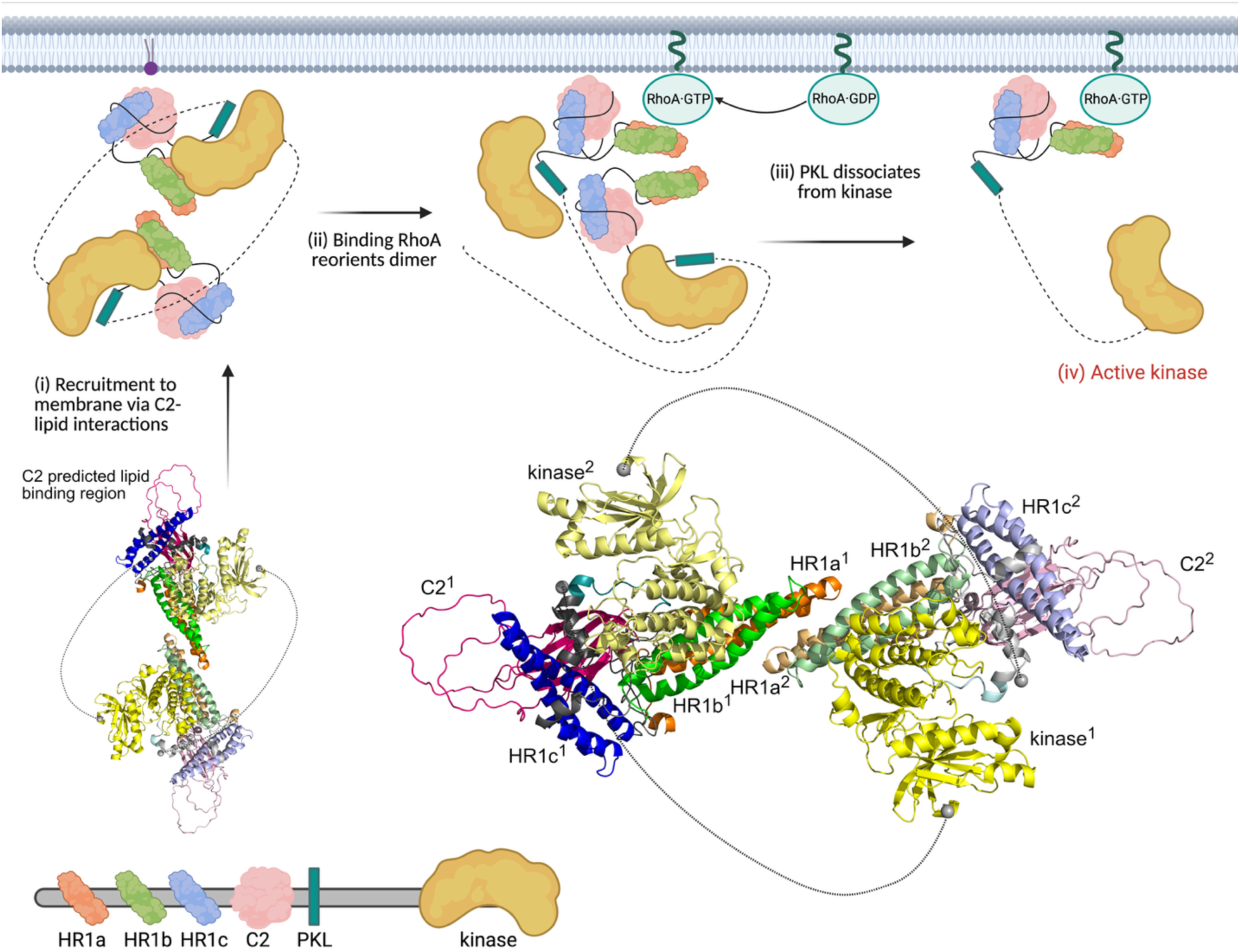
Model of the role of HR1a dimers in PKN1 activation. The model of full-length PKN1 was created using HR1abc dimer and Alphafold3 models. The two monomers are shown in darker and paler shades of the colours in the domain schematic (bottom). The long unstructured linkers between the PKL and kinase domain are shown as dotted lines between the edges of the long loop (grey spheres). The C2-like domain lipid binding site is on the surface of the compact dimer and is free to be recruited by interactions with membrane lipids (i). Recruitment to the membrane by binding lipid (purple) brings the HR1 domains into proximity with active RhoA, which binds to the HR1a domain, reorienting the PKN1 dimer (ii). This reorientation destabilizes the intermolecular PKL-kinase interaction (iii), leading to release of the kinase and its subsequent activation (iv). Created with BioRender.com.

It is known that PKN1 is activated by binding RhoA and Rac1 (15; 17; 52; 53). Here we have shown, using AUC, ITC and SEC-SAXS, that RhoA forms a 1:2 complex with a PKN1 HR1a dimer. The interaction is asymmetric, which would theoretically allow another HR1a domain to be assembled onto HR1a^2^, although higher order polymers were not observed in the AUC experiments (Figure 5B). Presumably, the initial binding of HR1a^1^ to RhoA represents one event observed in the ITC experiments (Figure 5C). Once HR1a^1^ is bound, its rearrangement in a high affinity RhoA complex favours the interaction with HR1a^2^, presumably driven by subtle reorientations in HR1a^1^. It has been suggested that RhoA activates the PKNs by binding to HR1a and reducing auto-inhibition by engaging a pseudosubstrate region within this domain (16). However, we have shown here that binding to RhoA causes a dramatic reorientation of the HR1a dimer. A RhoA complex with the same stoichiometry is formed by PKN1 HR1a, HR1ab and HR1abc, so it seems likely that RhoA will also induce formation of a reoriented dimer in the context of the full-length protein. This reorientation could play a critical role in the activation of PKN1, by disrupting the compact dimer and preventing the PKL-kinase interaction, therefore the dimer observed in the RhoA-bound form is likely to be compatible with activation. PKN1 is also activated by lipids binding to the C2-like domain. The model suggests that lipid binding to the canonical site on the C2-like domain would not disrupt the PKL-kinase inhibitory interactions (Figure 8). Instead, lipid activation may act via membrane recruitment, which would promote PKN1 binding to Rho proteins. Rac1 also binds to PKN1 and activates the kinase (17) but we show here it binds with a 1:1 stoichiometry. Rac1 would therefore disrupt HR1a-mediated oligomers, leading to monomeric PKN1 and activation of the kinase. Collectively, our SAXS results differentiate the structural complex formed by PKN1 and RhoA from that formed by PKN1 and Rac1. These different complexes are likely to have implications for PKN1 activation and/or could affect PKN1 localization.

### Experimental Procedures

#### Expression and production of recombinant proteins

PKN1 HR1a (1–106), PKN1 HR1b (122–199), PKN1 HR1ab (1–199), PKN3 HR1a (1–80), PKN3 HR1b (100–171) and PKN3 HR1ab (1–171) were expressed as described previously (37; 35). PKN1 HR1c (201–297), PKN1 HR1abc (1–297), PKN3 HR1c (169–263), PKN3 HR1abc (1–263), RhoA (F25N, Q63L, 1-189) and Rac1 (Q61L, 1-188) were amplified and cloned into pGEX-6P-1 (Cytiva). Proteins were expressed in *E. coli* BL21 (Invitrogen) or BL21-Rosetta2 (DE3) pLysS (Novagen) for 20 h at 20 °C, purified with glutathione-agarose beads (Merck) and cleaved with HRV 3C protease. PKN1 HR1b, PKN1 HR1c, PKN3 HR1c, RhoA and Rac1 were purified on a Superdex 75 16/60 column (Cytiva) and PKN1 HR1abc on a Superdex 200 16/60 column (Cytiva). PKN1 HR1a, PKN1 HR1ab, PKN3 HR1a and PKN3 HR1b were purified on a glutathione sepharose column (Cytiva) followed by Superdex 200 16/60 (PKN1 HR1ab), Superdex 75 16/60 (PKN3 HR1a and PKN3 HR1b), or ResourceQ (Cytiva) followed by Superdex 200 16/60 (PKN1 HR1a). PKN3 HR1ab and PKN3 HR1abc were purified on a Heparin HiPrep FF 16/10 column (Cytiva) and then a Superdex 200 16/60 column. Protein concentrations were determined by amino acid analysis (Protein and Nucleic Acid Chemistry Facility, Department of Biochemistry, University of Cambridge).

#### Nucleotide exchange

The nucleotide on RhoA and Rac1 was exchanged for GMPPNP (Merck) and the bound nucleotide analysed by HPLC as described previously (54).

#### NMR spectroscopy

NMR experiments and resonance assignments of the PKN1 HR1c domain have been described (41). NMR experiments were recorded on 1.6 mM ^15^N- or 1.2 mM ^15^N, ^13^C-labelled PKN1 HR1c in 20 mM sodium phosphate pH 7.3, 150 mM NaCl, 0.05% NaN_3_ and 10% D_2_O. Heteronuclear NOE (4.5 s saturation time) and ^15^N-separated NOESY (150 ms mixing time) experiments were recorded on a Bruker DRX500. ^15^N T_1_ and T_2_ experiments were recorded on a Bruker DRX500 as pseudo-3D experiments, with time delays of 0.01, 0.05, 0.10, 0.15, 0.25, 0.40, 0.50, 0.60 and 0.80 s for the T_1_ and 0.0144, 0.0288, 0.0432, 0.0576, 0.0720, 0.0864 and 0.1008 s for the T_2_. Heteronuclear NOE, T_1_ and T_2_ data were analysed as described previously (55). ^13^C-separated NOESY (100 ms mixing time) was recorded on a Bruker AV800. NMR data were processed using Azara (Wayne Boucher, University of Cambridge). NMR spectra were analysed using CCPN Analysis 2.4 (56).

#### Structure calculations

The predicted dihedral angles from TALOS-N (57) and the intensities of peaks in ^15^N-separated NOESY/^13^C-separated NOESY spectra were used to derive dihedral angle and distance restraints, respectively. Structure calculations were performed using ARIA 1.2 (58) interfaced to CNS version 1.2 (59). Structures were calculated during 8 iterations where the ambiguity of restraints was reduced. A total of 100 structures were calculated in the final iteration and the 80 lowest energy structures were water-refined.

#### NMR titrations

^15^N HSQCs were recorded on 0.3 mM ^15^N-labelled PKN1 HR1c in 50 mM Tris-HCl pH 7.5, 150 mM NaCl, 0.05% NaN_3_ and 10% D_2_O (plus 5 mM DTT in the HR1ab titrations). HR1a titrations were recorded on a Bruker DRX500 at molar ratios of 1:0, 1:0.25, 1:0.5, 1:1, 1:2 and 1:6. HR1b (molar ratios of 1:0, 1:0.25, 1:0.5, 1:1 and 1:6) and HR1ab titrations (molar ratios of 1:0, 1:0.25, 1:0.5, 1:1, 1:2 and 1:6) were recorded on a Bruker AV800. ^15^N HSQCs were recorded on ^15^N-labelled PKN1 HR1ab on a Bruker AV800 in the same buffer at concentrations of 0.357 mM, 0.2 mM, 0.1 mM, 0.05 mM and 0.015 mM. All experiments were recorded at 25 ᵒC.

#### Chemical shift mapping

Combined chemical shift changes, 𝛥𝛿, were calculated for residues whose resonances could be assigned using the following equation:

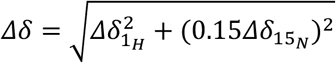

where 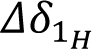 and 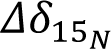 are the chemical shift changes in the ^1^H and ^15^N dimensions, respectively.

#### Circular dichroism

CD spectra were recorded on proteins at 0.2 mg/mL in 50 mM sodium phosphate pH 7.5, 150 mM NaF in a 0.1 mm pathlength quartz cuvette using an AVIV 410. The ellipticity was measured at 18 °C between 185 nm and 260 nm at 1 nm intervals with a 1 s averaging time/point. Four scans were recorded and averaged. For thermal melts, ellipticity at 222 nm was monitored at temperatures from 18 °C and 80 °C in 2 °C increments with a 30 s equilibration time. CD data were analysed using the DICHROWEB server and percentage helicity was estimated using the SELCON3 programme and reference set 3 (60; 61).

#### Analytical ultracentrifugation

Samples for AUC were loaded into double-sector Epon centrepieces with sapphire windows and a pathlength of 12 mm. These were loaded into an An60Ti rotor, equilibrated at 0 rpm for at least one hour and then run in an Optima XL-I centrifuge (Beckman Coulter) at 4 °C and 40,000 rpm. For the PKN1 HR1a-RhoA experiment, the proteins were incubated for 30 min before being loaded. Interference optics data were recorded in the continuous mode without averaging and with radial increments of 0.003 cm. Scans were recorded at 1 min 30 sec intervals until no further sedimentation was observed. The solvent density, solvent viscosity and the protein partial specific volume were calculated using SEDNTERP (62). A total of 200 scans were analysed and the data were fitted to a continuous c(s) distribution using a Marquardt-Levenberg algorithm in SEDFIT v14.1 (63). A Simplex algorithm was also used to check for consistent fit results. The c(s) model provided an estimate of the global frictional ratio, which is used to estimate the molecular weight of each species present. The figures were generated with the program GUSSI (64).

#### Small-angle X-ray scattering

SAXS data were collected on beamline B21 at the Diamond Light Source (65) at room temperature. SEC-SAXS was performed on: 1.5 mM HR1a; 1.5 mM HR1ab; 0.7 mM HR1abc; 0.41 mM RhoA + 0.41 mM HR1a; 0.30 mM RhoA + 0.30 mM HR1ab; 0.020 mM RhoA + 0.040 mM HR1abc; 0.33 mM Rac1 + 0.33 mM HR1a; and 0.25 mM Rac1 + 0.12 mM HR1ab. All GTPases were GMPPNP-bound. For each sample, 50 μL was injected onto a Superdex 200 increase 3.2/300 (Cytiva) and eluted in 50 mM Tris-HCl pH 7.5, 150 mM NaCl and 5 mM DTT (and 5 mM MgCl_2_ for any samples with a GTPase) at a flow rate of 0.075 mL/min.

Scattering curves were analysed with BioXTAS RAW v1.6.4 (66) using the ATSAS 2.8 package (67). Signal frames with a constant estimate of the radius of gyration, *R*_g_, were averaged and the buffer subtracted. PRIMUS (68) was used to perform Guinier analysis to determine *R*_g_ and estimate the molecular weight based on the volume of correlation, V_c_. The pair distance distribution function, p(r), was determined using GNOM (69) to obtain *D*_max_. A total of 15 *ab initio* models were generated in ‘slow mode’ by the program DAMMIF (70), aligned, averaged and filtered using the DAMAVER suite of programs (71) and further refined using DAMMIN (72), with a final χ^2^ value between 1.0 and 1.1. Resolution was estimated using SASRES (73).

#### Modelling

The PKN1 HR1a dimer was modelled from the SAXS data using FOXSDOCK (44; 45) and the free HR1a coordinates from the RhoA complex structure (PDB:1CXZ). The unstructured tails were built onto the dimer using Modeller (74) and then defined as flexible regions for MULTIFOXS (45) refinement. To assess sidechain interactions, FOXSDOCK and AlphaFold3 models were water-refined using the HADDOCK server refinement interface with the default parameters.

To construct a model of the HR1ab dimer, HADDOCK2.4 (47) was used to dock the HR1a + linker (residues 1-121) AlphaFold3 dimer with two HR1b monomers (PDB:2RMK). HR1b residues with chemical shift changes larger than the average plus one standard deviation between the free domain and in the context of HR1ab provided ambiguous interaction restraints for HR1b. For HR1a, all residues in the structured region (13–98) whose relative solvent-exposure was greater than 40% were used as passive residues. Interactions between HR1a and HR1b were only intramolecular, and restraints were applied to keep the dimer symmetric. A distance restraint was applied to keep the C-terminus of the linker close to the beginning of HR1b. All models in each HADDOCK cluster were fit to the SAXS data using FOXS. The cluster with the lowest average χ^2^ was selected and from this the model with the lowest χ^2^ value was chosen. The linker between HR1a and HR1b was replaced using Modeller10.5, the results were fit to the SAXS data and the model with the lowest χ^2^ value was selected.

A similar approach was used to model the HR1abc dimer. The chemical shift changes in HR1c when titrated with HR1ab were used to define the active residues in HR1c, with residues that shifted more than the average plus one standard deviation and more than 15% solvent exposed selected. For HR1ab, all residues that were more than 40% solvent exposed were defined as passive residues, with the exception of the long linker between HR1a and HR1b and the extended N- and C-termini.

To model the RhoA-HR1a dimer, a second HR1a monomer was docked onto the RhoA-HR1a complex structure (PDB:1CXZ, contact II) using FOXSDOCK. The flexible tails in the model with the lowest combined score (energy and χ^2^ score) were refined with MULTIFOXS, then water-refined using the HADDOCK server refinement interface with the default parameters.

HR1b domains were placed in random orientations with respect to the MULTIFOXS RhoA-HR1a dimer model and the loops built by Modeller. This was then refined with MULTIFOXS with the interdomain linker defined as flexible.

The Rac1-HR1b complex structure (PDB:2RMK) was used as a template to generate a model of Rac1-HR1a, which was refined using MULTIFOXS. The Rac1-HR1ab complex was built starting from the Rac1-HR1a model or the Rac1-HR1b structure, with the other HR1 domain added in a random orientation and the linker built with Modeller. The flexible termini and the linker were then refined using MULTIFOXS.

CRYSOL was used to calculate the scattering expected from a given structure and fit it to the experimental SAXS curve (75).

#### Isothermal titration calorimetry

ITC experiments were carried out in 50 mM Tris-HCl pH 7.5, 150 mM NaCl, 5 mM MgCl_2_, 5 mM DTT using an iTC_200_ (Cytiva) at 25 °C. 1 mM RhoA·GMPPNP was injected into 100 μM HR1a. The data were integrated using NITPIC and then fitted with SEDPHAT, using the Simplex algorithm (76). The figure was prepared with GUSSI (64).

## Data Availability

NMR structure of PKN1 HR1c has been deposited in the protein data bank (9RXE). Other models are freely available from the corresponding authors on request. All other data is contained within the manuscript.

## Supporting Information

This article contains supporting information (50).

## Acknowledgements

We are grateful to the Biophysics Facility in the Department of Biochemistry and Dr Katherine Stott for her help with ITC and AUC data analysis.

## Funding and additional information

This work was supported by the A.G. Leventis Foundation (to G.S.) and the Medical Research Council (to D.O. and H.R.M.) award number MR/J007803/1.

