## Supplementary material for "Structural basis of the protein kinase PKN1 HR1 domain oligomerization and differential regulation by RhoA and Rac1": Supplemetary Figures 1-11; Supplementary Tables 1A, 1B, 2

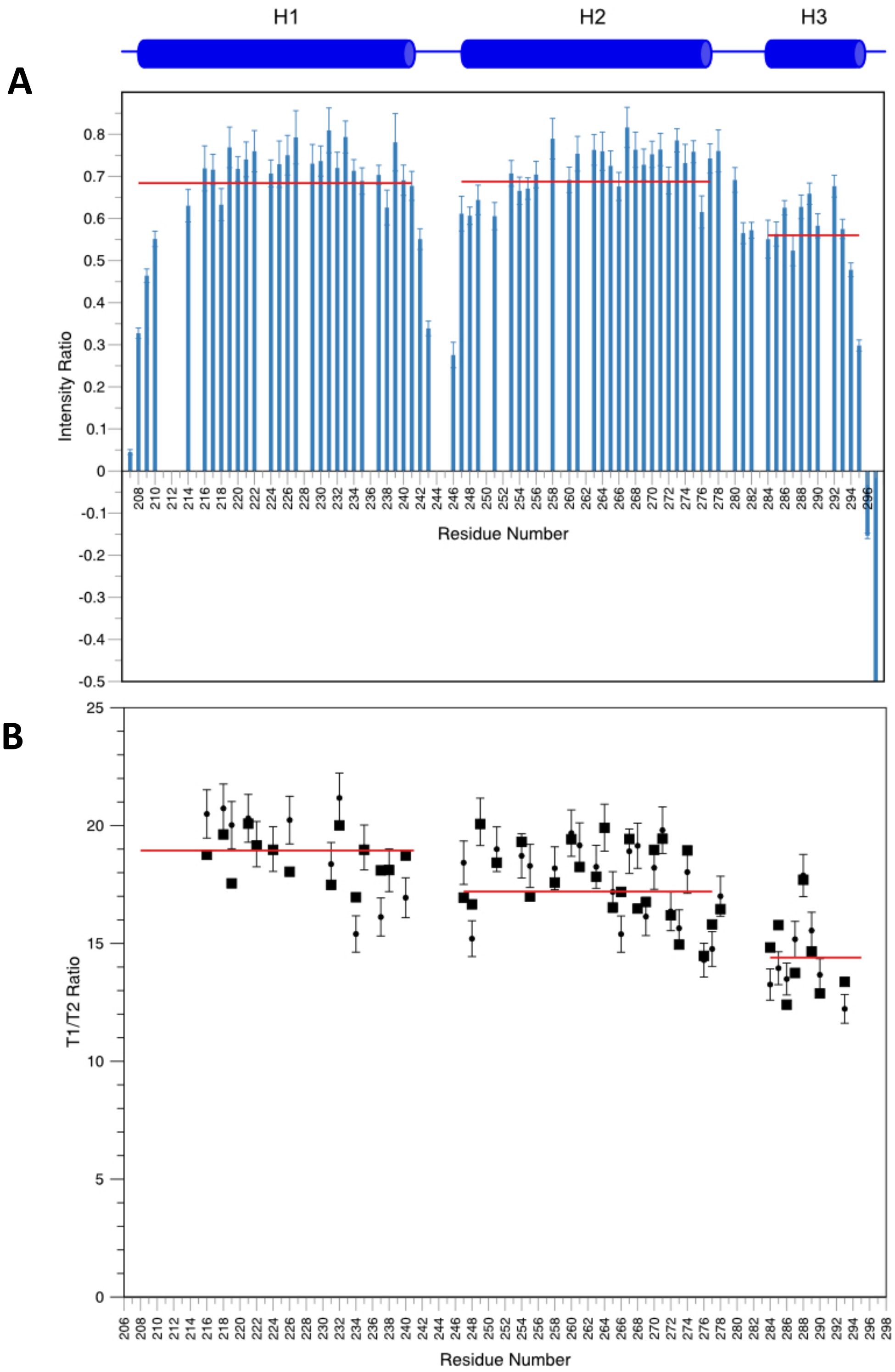


**Supplementary Figure 1. $^{15}$N relaxation data of PKN1 HR1ab.** (A) $^{1}$H-$^{15}$N heteronuclear NOE intensity ratios. The average NOE for each individual helix is indicated by a solid line. (B) Experimental T1/T2 ratios were used to calculate the rotational diffusion tensor. The experimental values are shown as small black circles with their associated errors and the average T1/T2 across each helix is indicated by a solid line. The calculated T1/T2 ratios from the diffusion tensor are shown as large squares.

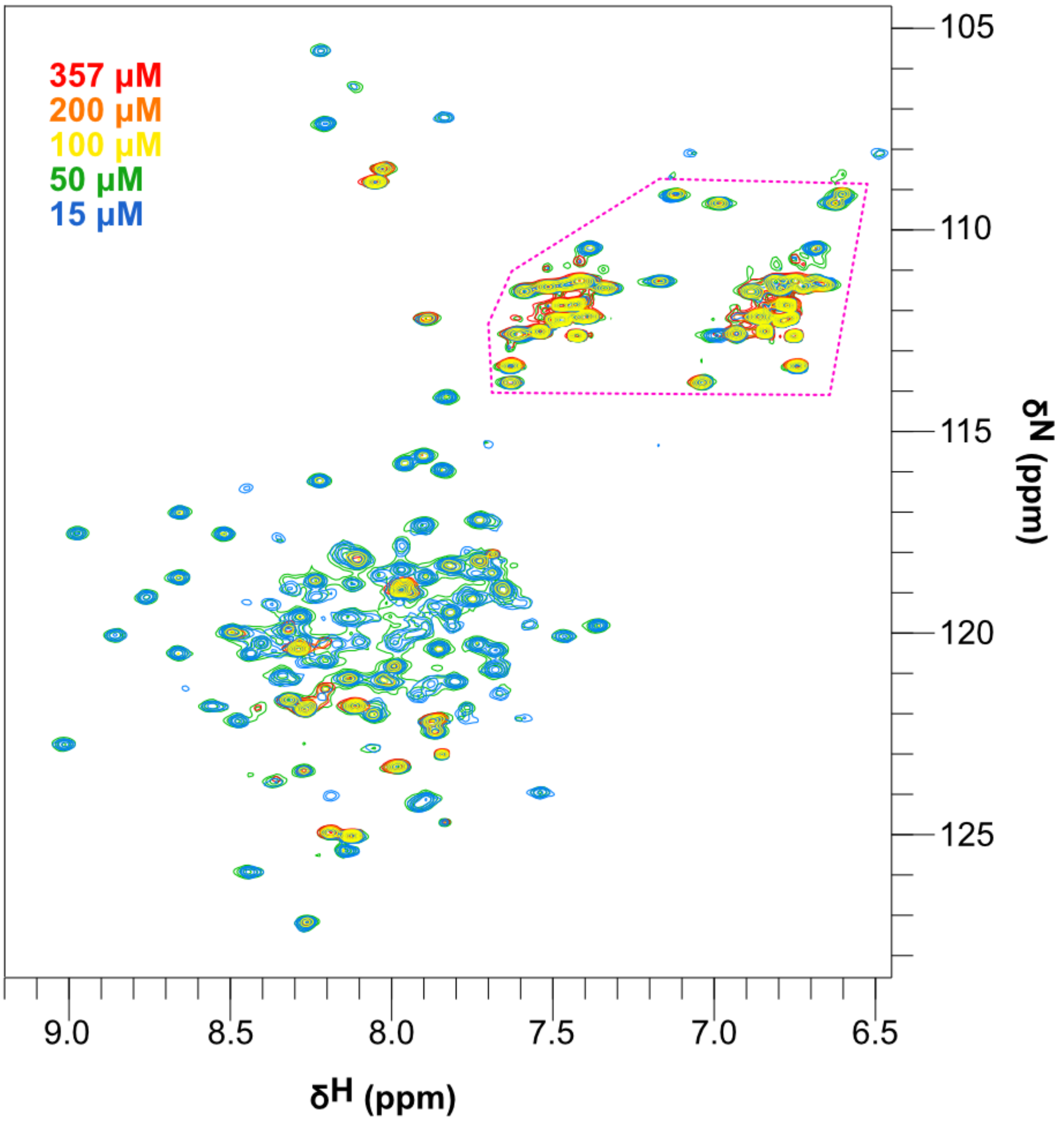


**Supplementary Figure 2. $^{15}$N-HSQC spectrum of PKN1 HR1ab.** $^{15}$N-HSQC spectra of PKN1 HR1ab in 50 mM Tris-HCl pH 7.5, 150 mM NaCl, 5 mM DTT and 10% $D_2O$ at concentrations ranging from 357 μM to 15 μM coloured as shown in the key on the top left. The region encompassed by the pink dashed line shows several peaks from sidechain $NH_2$ groups. These, and a few other amides are visible in all the spectra, but the majority of the peaks are only apparent at the two lowest concentrations. The peak intensities were normalised to Ala198, which is visible at all concentrations.

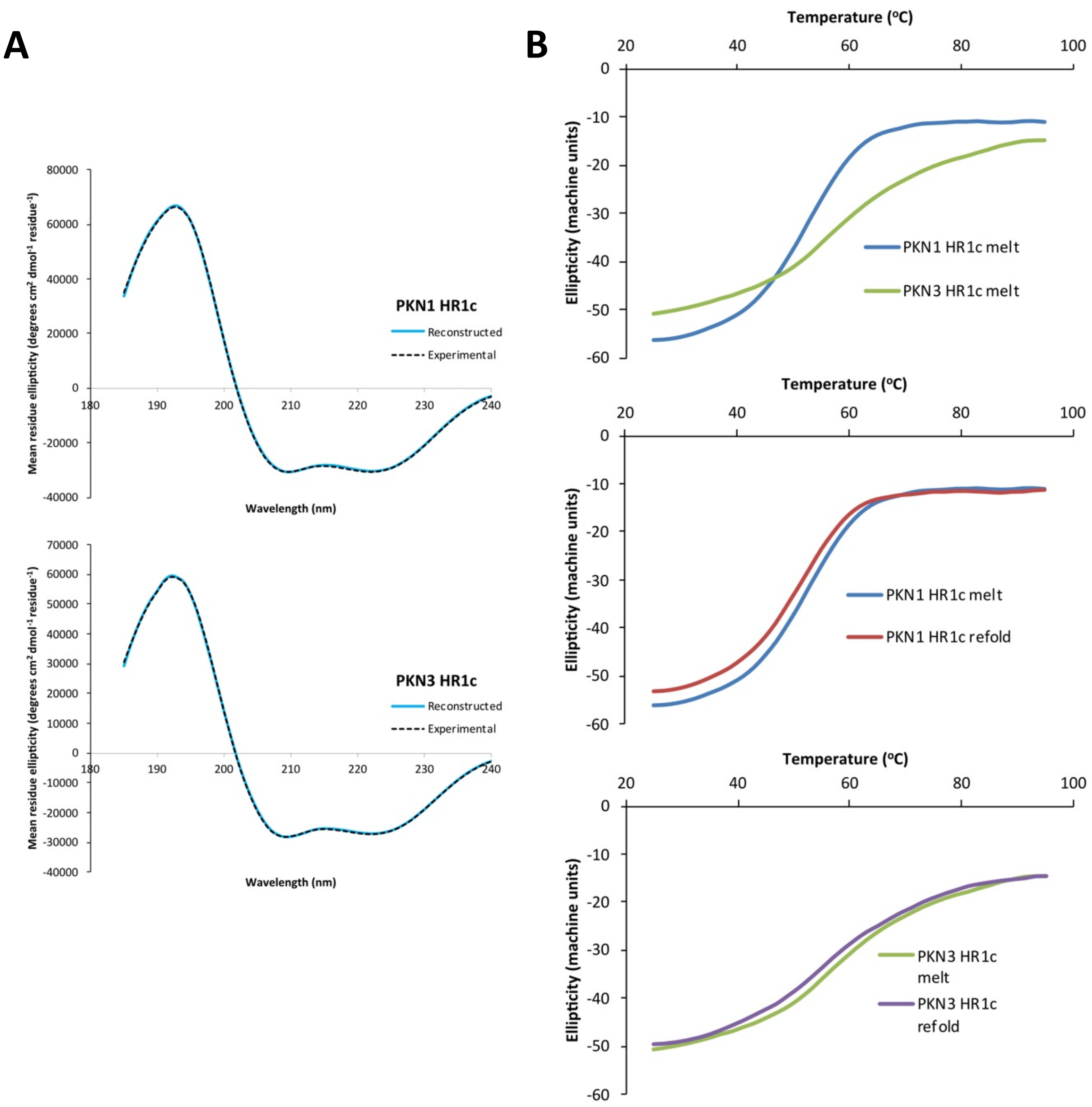


**Supplementary Figure 3**. Comparison of PKN1 HR1c and PKN3 HR1c. (A) CD Spectra: PKN1 HR1c and PKN3 HR1c wavelength scans recorded with 0.2 mg/mL protein. The experimental data is overlaid with the reconstructed data from the DICHROWEB analysis. (B) Thermal stability: the thermal melt was obtained by recording the ellipticity at 222 nm with increasing temperature. Top panel: a comparison of the melt curves of PKN1 and PKN3 HR1c domains. The lower two panels show the ellipticity at 222 nm, of each domain separately, as the temperature was increased (melt) and then reduced back to the starting temperature (refold).

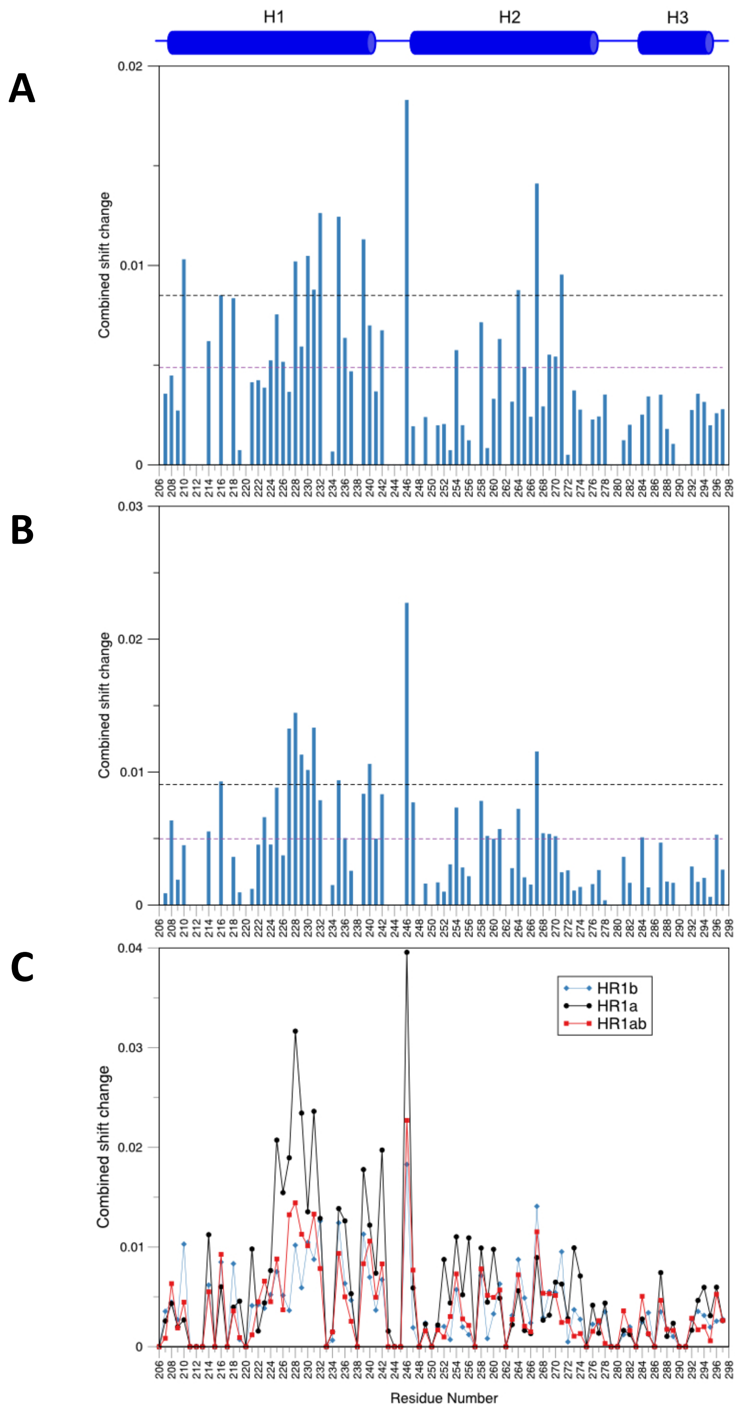


**Supplementary Figure 4. HR1c interactions with HR1a, HR1b and HR1ab.** Chemical shift changes (at 1:6 ratio) for backbone NH groups of residues 206-297 are shown. Residues which could not be reliably assigned due to overlap were assigned a combined shift change, Δδ, value of zero. (A) HR1c + HR1b chemical shift changes (B) HR1c + HR1ab chemical shift changes. The mean Δδ and mean Δδ + one standard deviation are marked with purple and black, dotted lines, respectively. (C) Comparison of shift changes when HR1c was titrated with HR1a, HR1b or HR1ab.

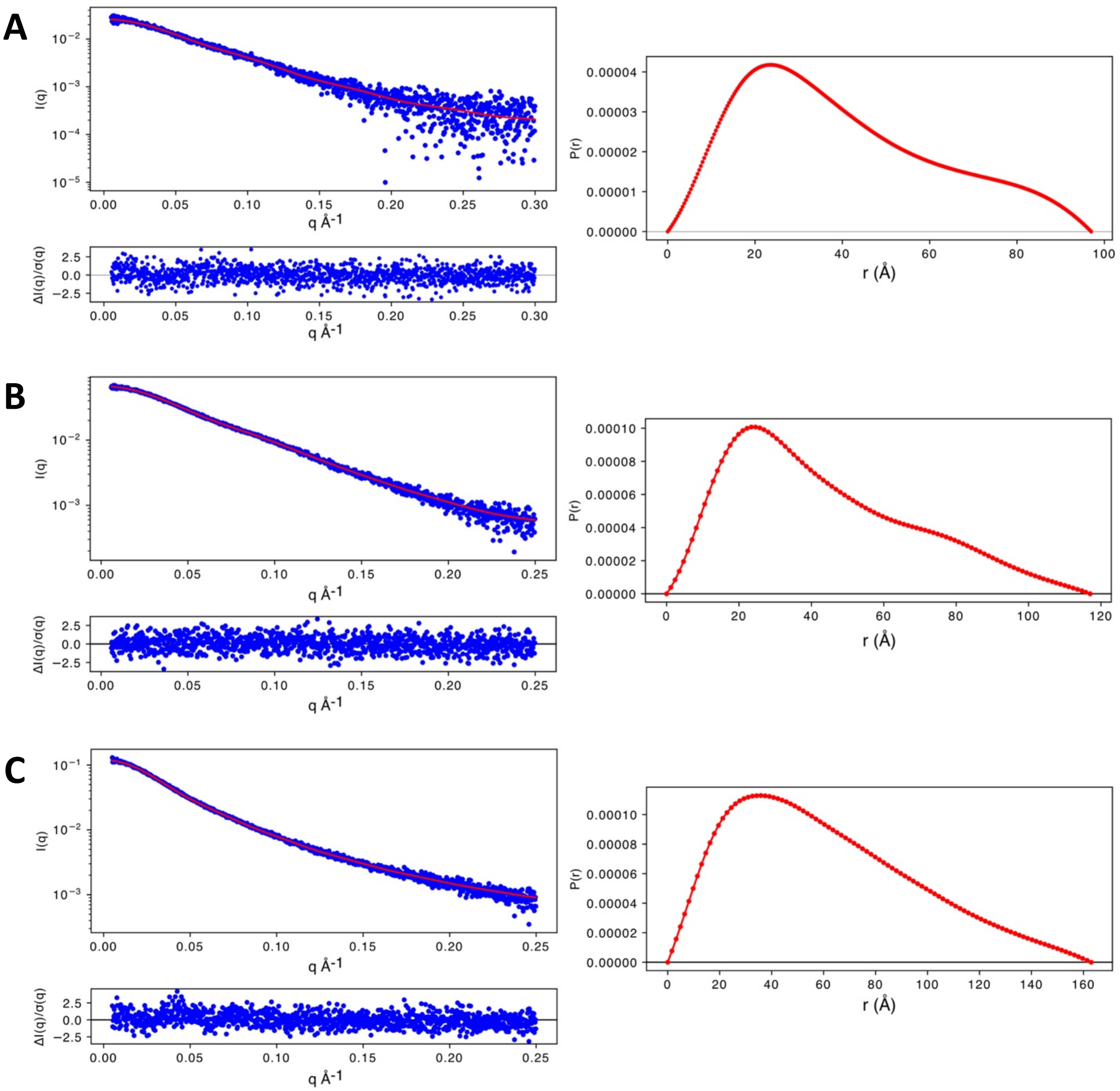


**Supplementary Figure 5. SAXS data analysis of HR1 dimers.** Scattering profiles and p(r) plots of (A) HR1a (B) HR1ab (C) HR1abc constructs from PKN1. Left panels: Scattering profile (data in blue). The red line represents the ATSAS fit to the data from which the $R_g$, MW estimation and p(r) plots were derived. Right panel: p(r) plot.

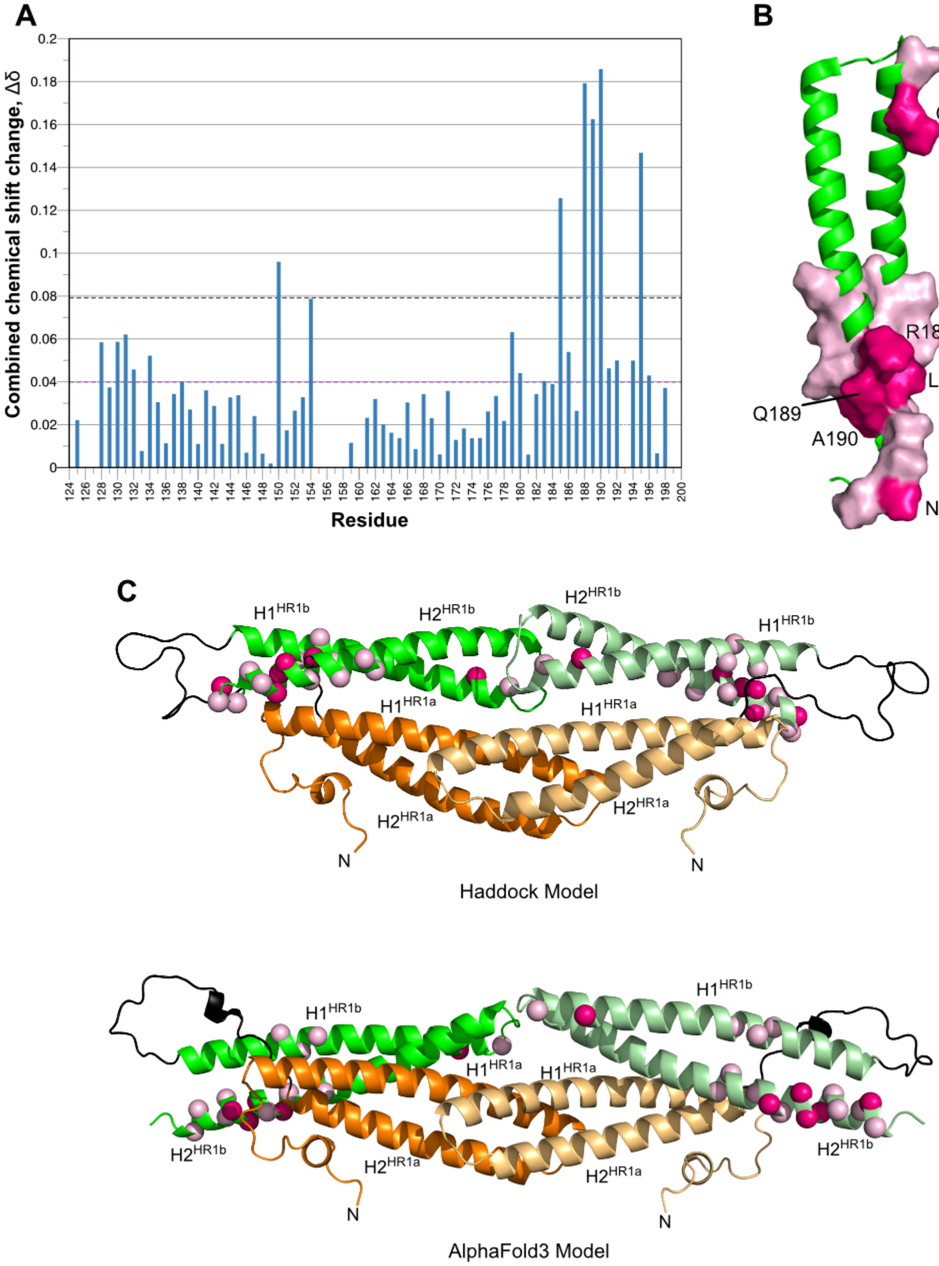


**Supplementary Figure 6. Chemical shifts of free HR1b vs. HR1b in the context of the HR1ab construct.** (A) The mean Δδ and mean Δδ + 1 SD are marked with purple and black, dotted lines, respectively. (B) HR1b residues whose resonances shifted significantly (mean + 1 SD) are shown in a surface representation in dark pink on the structure, while residues whose resonances shifted only above the mean are coloured on the HR1b structure in pale pink. (C) Comparison of the HR1ab HADDOCK-derived model (above) and the AlphaFold3 model (below) that best fits the SAXS data. HR1a is shown in orange, HR1b is green and the linker between them is black. Residues in HR1b affected by the presence of HR1a are coloured as in B.

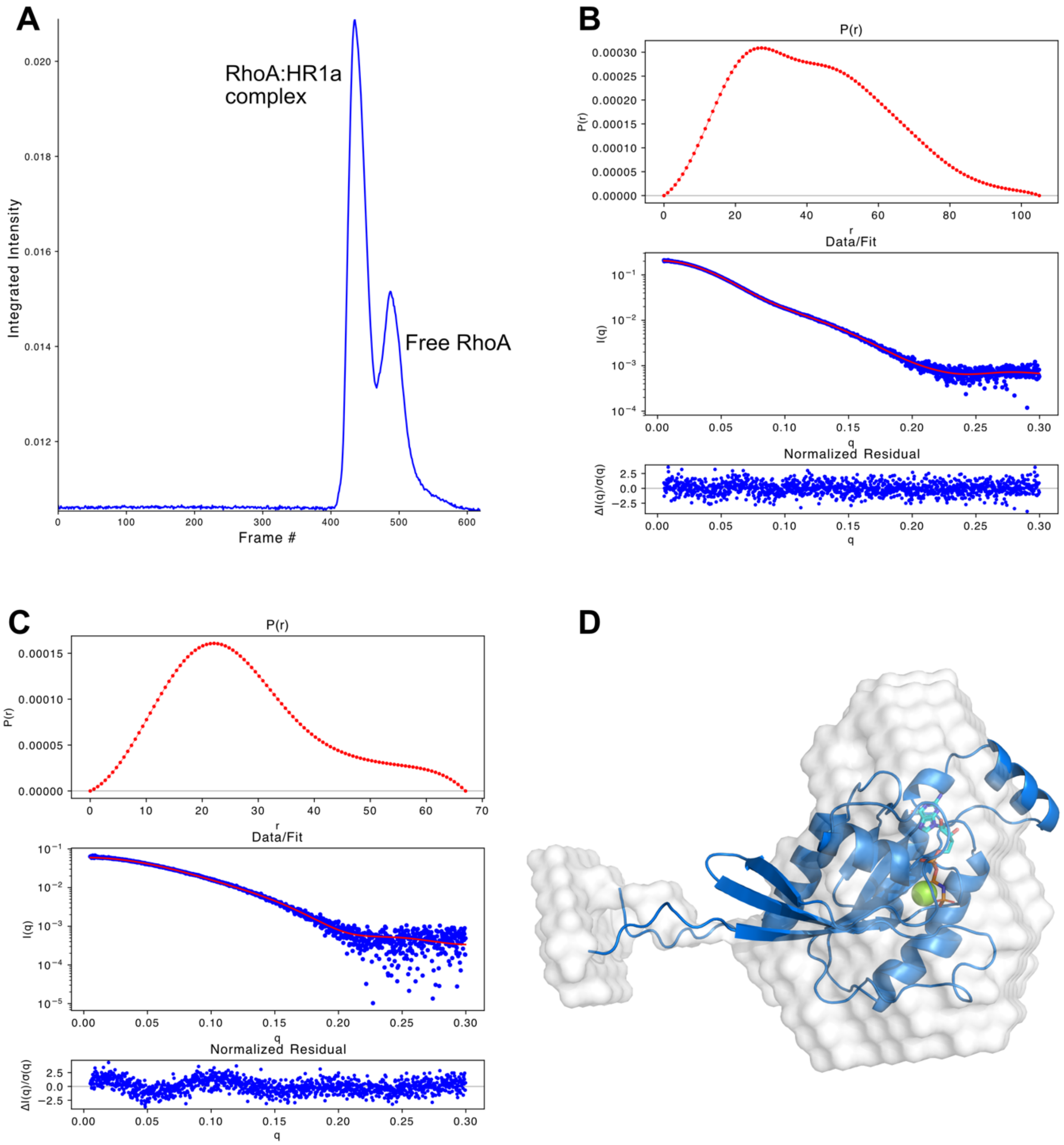


**Supplementary Figure 7. SAXS data analysis of the RhoA/HR1a complex.** (A) A representative chromatogram from SEC-SAXS obtained when RhoA was mixed with HR1a (or HR1ab) in equimolar ratios. Since RhoA forms 1:2 complexes with HR1a-containing constructs, the second peak is assumed to correspond to excess RhoA. (B,C) Scattering profiles and p(r) plots of (B) RhoA/HR1a (C) free RhoA. Upper panels: p(r) plots. Middle panels: Scattering profiles. The data are shown in blue. The red lines represent the ATSAS fit to the data from which the $R_g$, MW estimation and p(r) plots were derived. Bottom panels: the residuals from the ATSAS fit. (D) Low resolution *ab initio* model of RhoA overlaid with the best MULTIFOXS model of RhoA.

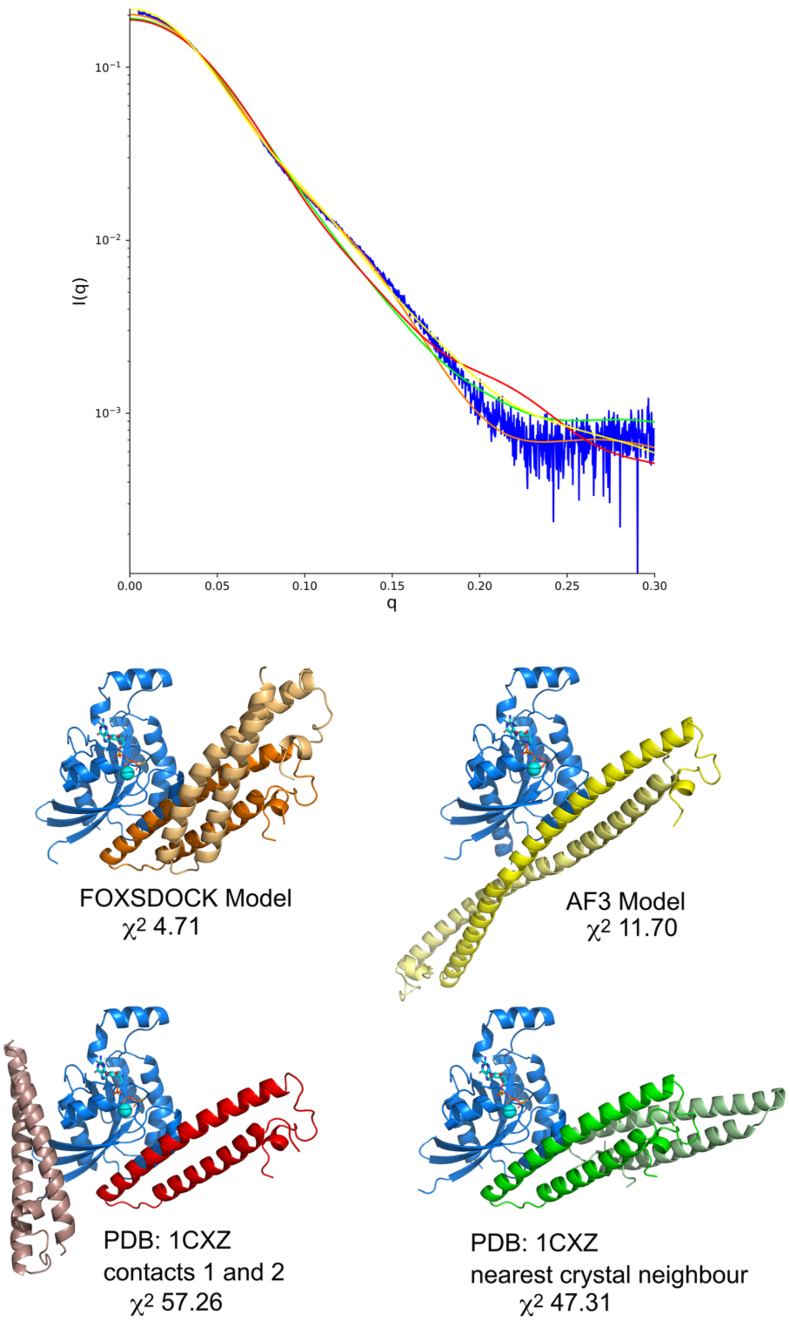


**Supplementary Figure 8. Solution scattering evaluation of possible RhoA:HR1a 1:2 models and fitting to experimental SAXS data using CRYSOL.** The SAXS data is shown in blue, with the fits of the different models shown below coloured as follows: the FOXSDOCK model (orange), RhoA with a monomeric HR1a domain bound at each of the two contact sites identified in the X-ray structure (red, PDB: 1CXZ), RhoA bound to an HR1a dimer predicted by AlphaFold3 (yellow), RhoA bound to an HR1a dimer formed by HR1a interacting with its nearest neighbour in the crystal (green, generated from PDBePISA). These models and the FOXSDOCK model were analysed with CRYSOL and the $\chi^2$ value is indicated next to each model.

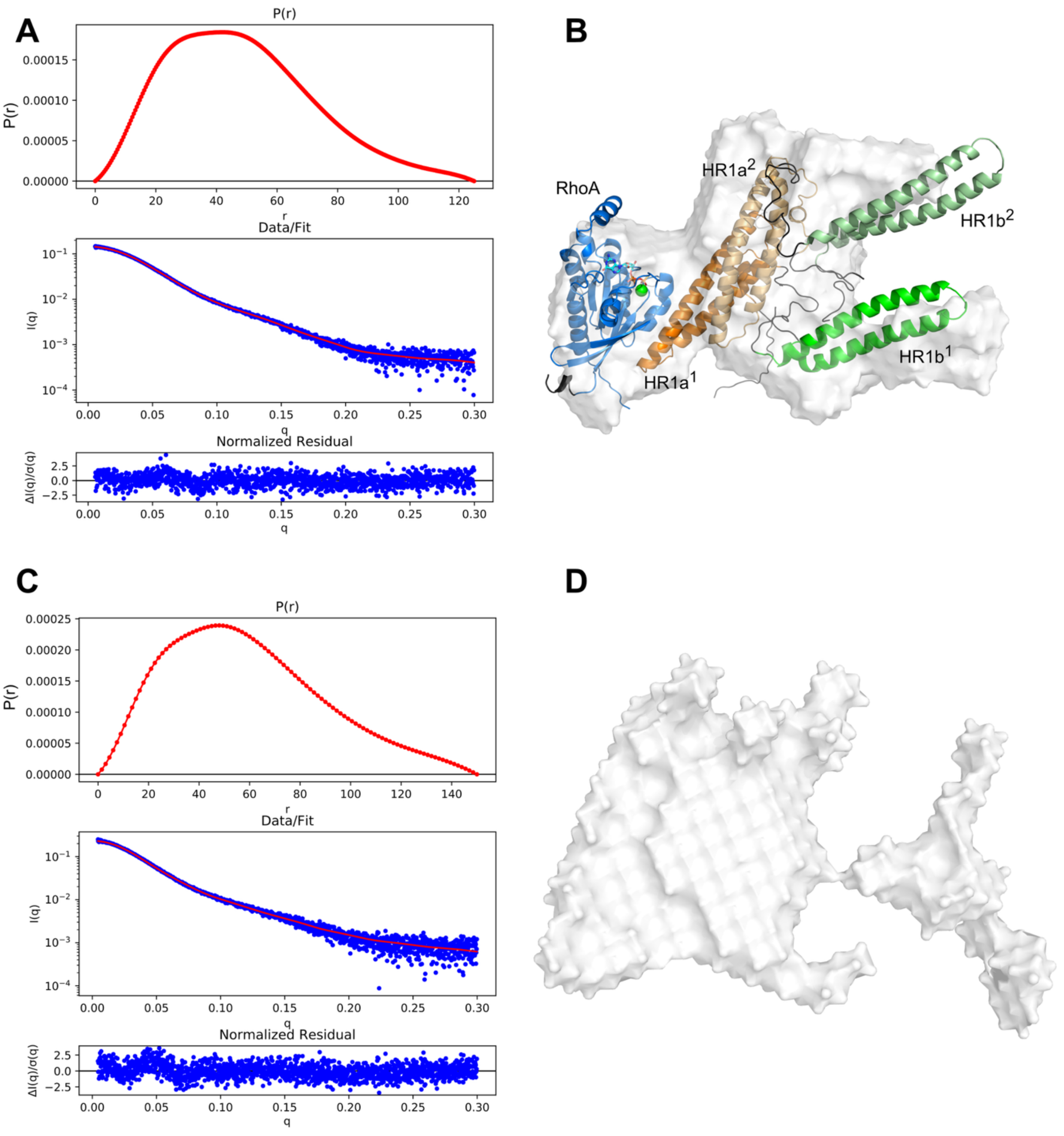


**Supplementary Figure 9. SAXS data analysis of RhoA in complex with HR1ab and HR1abc.** Scattering profiles and p(r) plots of (A) RhoA/HR1ab (C) RhoA/HR1abc. Upper panels: p(r) plots. Middle panels: Scattering profiles. The data are shown in blue. The red lines represent the ATSAS fit to the data from which the $R_g$, MW estimation and p(r) plots were derived. Bottom panels: the residuals from the ATSAS fit. (B) Low resolution *ab initio* model (white) of RhoA/HR1ab overlaid with the best MULTIFOXS model of RhoA/HR1ab. RhoA is blue, with the nucleotide shown in stick representation. HR1a is orange and HR1b is green, with $HR1a^1$ and $HR1b^1$ in darker shades. (D) Low resolution *ab initio* model of RhoA/HR1abc.

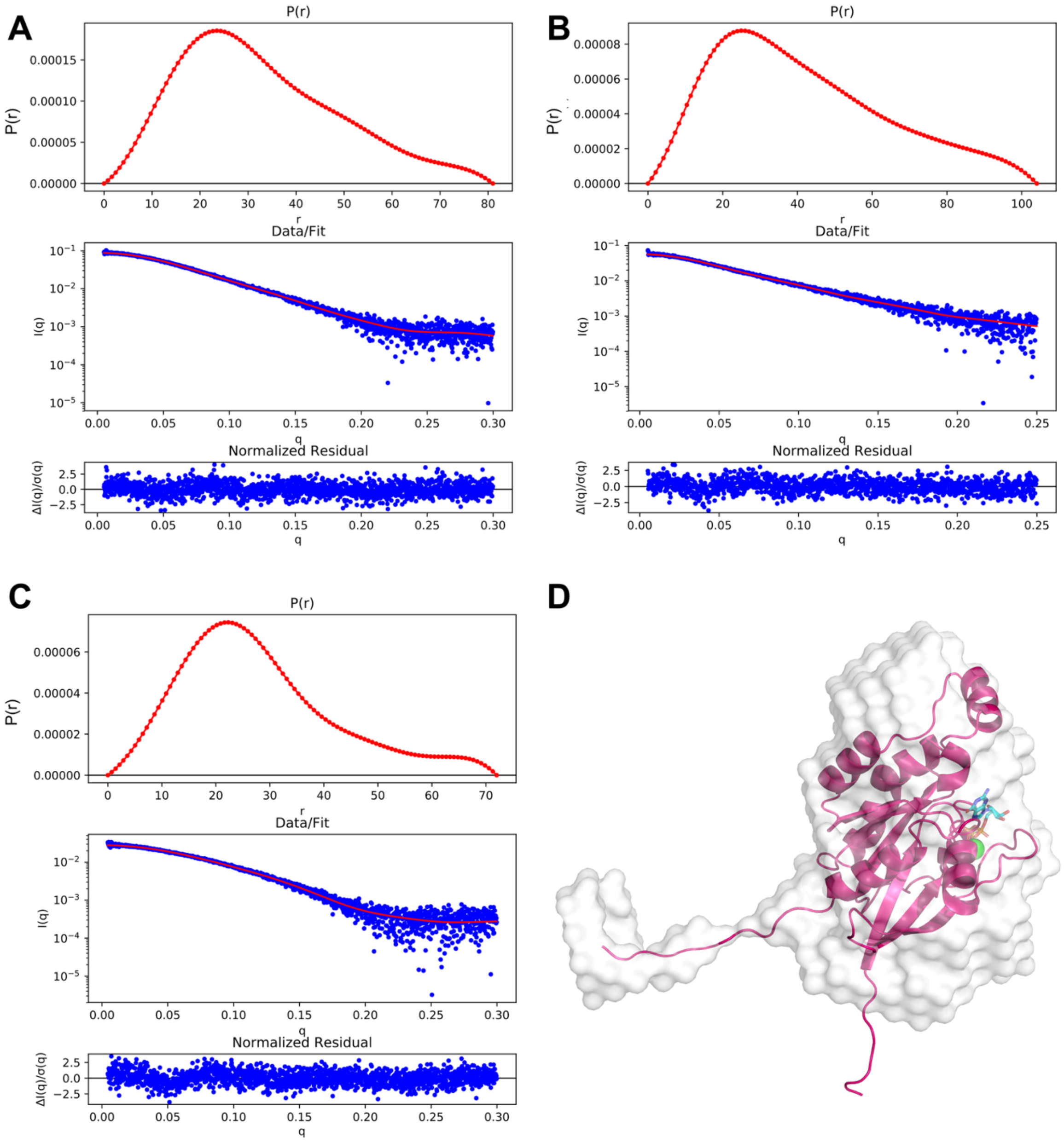


**Supplementary Figure 10. SAXS data analysis of Rac1 in complex with HR1 constructs.** Scattering profiles and p(r) plots of (A) Rac1-HR1a, (B) Rac1-HR1ab (C) Rac1 (excess in Rac1-HR1ab mixture). Upper panels: p(r) plots. Middle panels: Scattering profiles. The data are shown in blue. The red lines represent the ATSAS fit to the data from which the $R_g$, MW estimation and p(r) plots were derived. Bottom panels: the residuals from the ATSAS fit. (D) Low resolution *ab initio* model of Rac1 shown as a white surface overlaid with the best MULTIFOXS model of Rac1. Rac1 is shown in pink.

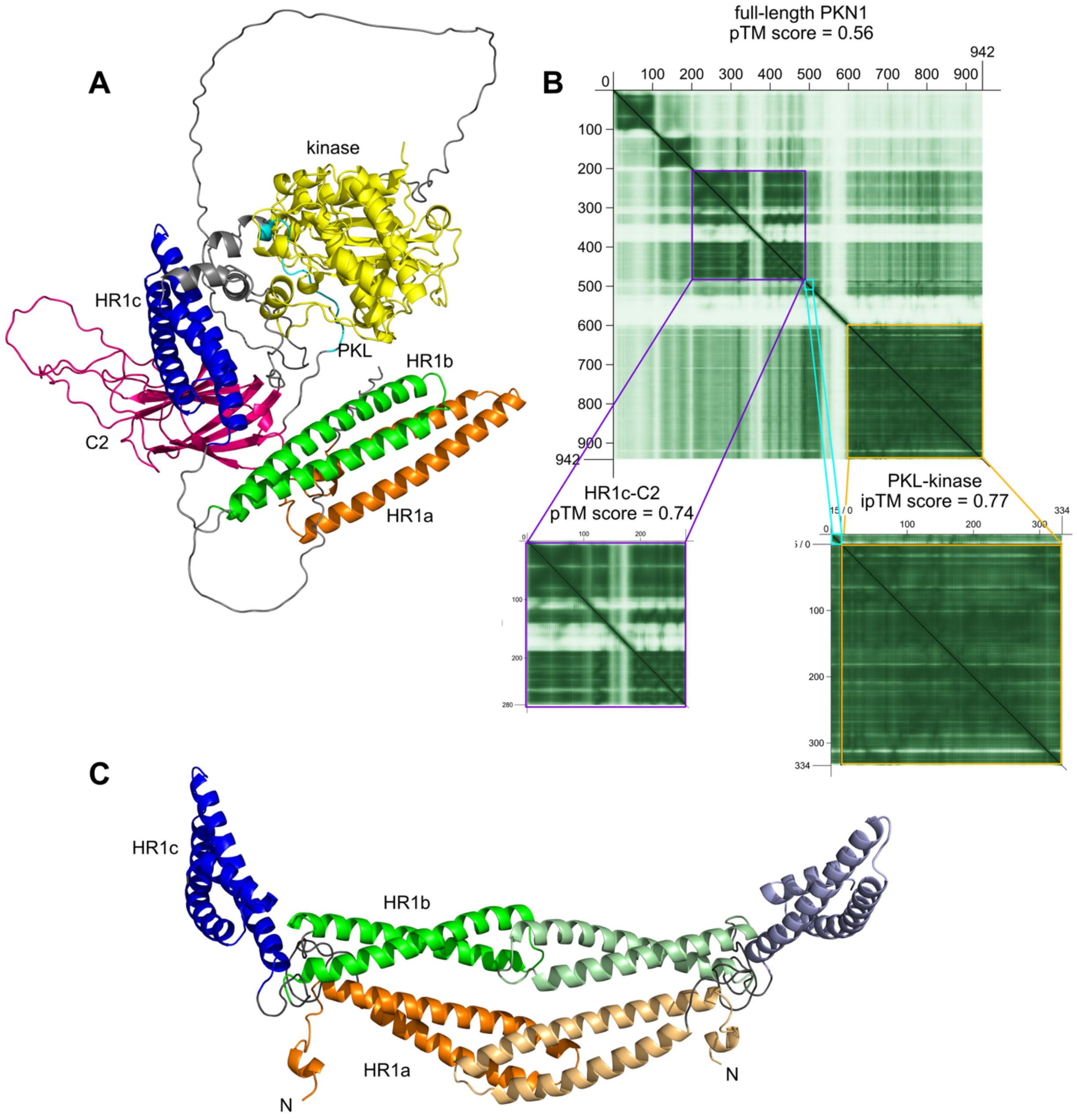


**Supplementary Figure 11. AlphaFold3 model of PKN1.** (A) Full-length PKN1 model with the domains and PKL region labelled. (B) Predicted align error (PAE) plots and scores of full-length PKN1 (main panel), where darker off-diagonal colours indicate higher confidence in interdomain interactions. Lower panels show the scores and PAE plots of the HR1c-C2 prediction (left) and the PKL-kinase complex prediction (right). PAE plots were generated using PAE viewer (50). (C) The HR1abc dimer model, rotated so that the HR1c domain is at the same orientation as HR1c in Panel A.

**Supplementary Table 1A. PKN1 AUC Data Analysis.**

| Sample | Protein concentration (μM) | Unnormalised RMSD of the fit | Species detected |
|---|---|---|---|
| PKN1 HR1a | 100 | 0.0268 | 59.6% monomer: *s*=0.974<br>40.4% dimer: *s*=1.64 |
| | 200 | 0.0485 | 41.4% monomer: *s*=1.04<br>58.6% dimer: *s*=1.77 |
| | 400 | 0.1069 | 26.5% monomer *s*=1.15<br>73.5% dimer *s*=1.79 |
| PKN1 HR1b<br>Calculated 10.7 kDa<br>(Expected 9.1kDa) | 200 | 0.0284 | Monomer: *s*=0.737 |
| | 400 | 0.0402 | Monomer: *s*=0.726 |
| | 800 | 0.0349 | Monomer: *s*=0.729 |
| PKN1 HR1c<br>Calculated 13.2 kDa<br>(Expected 11.3 kDa) | 400 | 0.0372 | Monomer: *s*=0.768 |
| | 800 | 0.0137 | Monomer: *s*=0.773 |
| PKN1 HR1ab | 100 | 0.0266 | 56.6% monomer: *s*=1.28<br>43.4% dimer: *s*=2.29 |
| | 200 | 0.0775 | 39.1% monomer: *s*=1.32<br>60.9% dimer: *s*=2.28 |
| | 400 | 0.2030 | 22.5% monomer: *s*=1.33<br>77.5% dimer: *s*=2.23 |
| PKN1 HR1abc | 100 | 0.0330 | 47.8% monomer: *s*=1.36<br>23.2% dimer: *s*=2.02<br>29.0% trimer: *s*=2.74 |
| | 400 | 0.2514 | 9.1% monomer: *s*=1.30<br>18.2% dimer: *s*=1.69<br>72.7% trimer: *s*=2.53 |
| | 500 | 0.3512 | 10.7% monomer: *s*=1.38<br>10.8% dimer: *s*=1.75<br>78.5% trimer: *s*=2.52 |
| **RhoA Complex** | | | |
| RhoA | 150 | 0.0210 | RhoA: *s*=1.53 |
| RhoA + PKN1 HR1a | 150/150 | 0.0410 | 22.8% RhoA: *s*=1.54<br>77.2% RhoA-HR1a: *s*=2.64 |

**Supplementary Table 1B. PKN3 AUC Data Analysis.**

| Sample | Protein concentration (μM) | RMSD of the fit | Species detected |
|---|---|---|---|
| PKN3 HR1a | 100 | 0.0137 | 89.9% monomer: *s*=0.782<br>10.1% dimer: *s*=1.49 |
| | 200 | 0.0302 | 78.1% monomer: *s*=0.713<br>21.9% dimer: *s*=1.15 |
| | 400 | 0.0409 | 71.9% monomer: *s*=0.729<br>28.1% dimer: *s*=1.15 |
| PKN3 HR1b | 200 | 0.0224 | 90.3% monomer: *s*=0.716<br>9.7% dimer: *s*=1.19 |
| | 400 | 0.0313 | 85.1% monomer: *s*=0.727<br>14.9% dimer: *s*=1.26 |
| PKN3 HR1c | 100 | 0.0181 | 49.1% monomer: *s*=0.666<br>44.7% dimer: *s*=0.949<br>6.2% tetramer: *s*=1.68 |
| | 200 | 0.0141 | 36.4% monomer: *s*=0.606<br>56.8% dimer: *s*=1.02<br>6.8% tetramer: *s*=1.63 |
| | 400 | 0.0279 | 34.7% monomer: *s*=0.731<br>49.9% dimer: *s*=1.10<br>15.4% tetramer: *s*=1.51 |
| PKN3 HR1ab | 50 | 0.0207 | 91.4% monomer: *s*=1.08<br>8.6% dimer: *s*=1.55 |
| PKN3 HR1abc | 50 | 0.0363 | 13.5% monomer: *s*=1.42<br>22.4% dimer: *s*=2.10<br>55.8% trimer: *s*=2.78<br>8.3% higher oligomer: *s*=4.04 |
| | 100 | 0.0443 | 12.9% monomer: *s*=1.51<br>19.1% dimer: *s*=2.34<br>49.5% trimer: *s*=2.89<br>18.5% higher oligomer: *s*=4.13 |
| | 200 | 0.0655 | 5.1% monomer: *s*=1.47<br>14.1% dimer: *s*=2.23<br>54.5% trimer: *s*=3.01<br>26.3% higher oligomer: *s*=4.51 |

**Supplementary Table 2. SAXS Data Analysis.**

| Sample | Guinier analysis | | | p(r) analysis | | | Envelope | | | Model | | |
|---|---|---|---|---|---|---|---|---|---|---|---|---|
| | $R_g$ (Å) [a] | Molecular Mass (kDa) | | $R_g$ (Å) [a] | $D_{max}$ (Å) [b] | $\chi^2$ | Resolution (Å) | $\chi^2$ | Normalised Spatial Discrepancy | $R_g$ (Å) [a] | $D_{max}$ (Å) [b] | $\chi^2$ |
| | | Volume of Correlation | Theoretical | | | | | | | | | |
| HR1a | 32.56 | 29.0 | 24.65 | 31.76 | 97 | 1.064 | 34±3 | 1.022 | 1.142±0.036 | 28.50 (DOCK)<br>30.02 (AF3) | 101 (DOCK)<br>96 (AF3) | 1.05 (DOCK)<br>1.01 (AF3) |
| HR1ab | 33.48 | 36.7 | 44.68 | 35.10 | 117 | 1.063 | 34±3 | 1.016 | 1.189±0.035 | 33.4 | 120 | 1.51 |
| HR1abc | 47.48 | 67.4 | 66.28 | 49.13 | 163 | 1.049 | 51±4 | 1.043 | 1.318±0.071 | 47.6 | 168 | 1.67 |
| RhoA + HR1a | 32.27 | 55.6 | 46.36 | 32.57 | 105 | 1.069 | 37±3 | 1.096 | 1.312±0.058 | 29.18 | 103 | 2.23 |
| RhoA + HR1ab | 37.42 | 79.4 | 73.66 | 37.97 | 125 | 1.084 | 41±3 | 1.069 | 1.436±0.063 | 37.79 | 135 | 1.29 |
| RhoA+ HR1abc | 46.15 | 101.7 | 87.99 | 46.77 | 150 | 1.192 | 45±3 | 1.098 | 1.456±0.095 | | | |
| Rac1 + HR1a | 25.54 | 28.6 | 33.87 | 25.71 | 81 | 1.056 | 32±3 | 1.085 | 0.864±0.126 | 25.92 | 78 | 1.02 |
| Rac1 + HR1ab | 33.21 | 38.6 | 43.88 | 33.34 | 104 | 1.042 | 37±3 | 1.003 | 1.563±0.110 | 34.20 [c]<br>32.91 [d] | 115<br>115 | 1.04<br>1.18 |
| RhoA$_{HR1a}$ | 22.12 | 22.8 | 21.7 | 21.55 | 67 | 1.435 | 19±2 | 1.416 | 0.618±0.137 | 17.58 | 70.6 | 3.46 |
| RhoA$_{HR1ab}$ | 21.70 | 21.5 | 21.7 | 21.53 | 70 | 1.182 | 18±2 | 1.178 | 0.538±0.012 | 17.35 | 68 | 2.65 |
| Rac1$_{HR1ab}$ | 22.18 | 21.3 | 21.54 | 22.08 | 72 | 1.145 | 24±2 | 1.137 | 0.805±0.164 | 18.61 | 71 | 1.54 |

[a] Radius of gyration.
[b] Approximate maximum dimension of model; when more than one model is predicted, $D_{max}$ of each model is shown.
[c] Calculated with HR1a contacting Rac1.
[d] Calculated with HR1b contacting Rac1.
